# Mechanotransduction-dependent control of stereocilia dimensions and row identity in inner hair cells

**DOI:** 10.1101/731455

**Authors:** Jocelyn F. Krey, Paroma Chatterjee, Rachel A. Dumont, Dongseok Choi, Jonathan E. Bird, Peter G. Barr-Gillespie

## Abstract

Actin-rich structures like stereocilia and microvilli are assembled with precise control of length, diameter, and relative spacing. We found that developmental widening of the second-tallest stereocilia rank (row 2) of mouse inner hair cells correlated with the appearance of mechanotransduction. Correspondingly, *Tmc1*^*KO/KO*^;*Tmc2*^*KO/KO*^ or *Tmie*^*KO/KO*^ hair cells, which lack transduction, have significantly altered stereocilia lengths and diameters. EPS8 and the short splice isoform of MYO15A, identity markers for row 1 (tallest), lost their row exclusivity in transduction mutants, a result that was mimicked by block of transduction channels. Likewise, the heterodimeric capping protein subunit CAPZB and its partner TWF2 lost their row 2 tip localization in mutants, and GNAI3 failed to accumulate at row 1 tips. Redistribution of marker proteins was accompanied by increased variability in stereocilia height. Transduction channels thus specify and maintain row identity and control addition of new actin filaments to increase stereocilia diameter.

## Introduction

The vertebrate hair bundle relies on the precise assembly of its structure to carry out mechanoelectrical transduction, conversion of mechanical stimuli into cellular electrical responses [1]. The bundle consists of about 100 actin-filled stereocilia, which are arranged in rows of increasing height and form a staircase [2]. During development, growth of stereocilia follows a choreographed process whereby lengthening and widening are activated at discrete times [3]. While more thoroughly characterized in chick cochlea, stereocilia growth in mammalian cochlear hair cells follows a similar process, although lengthening and widening steps appear to be less clearly distinguished [4, 5].

Hair bundles of inner hair cells (IHCs) in the mammalian cochlea have only three rows of stereocilia, each of which are unique are in their dimensions: row 1 is tall and thick, row 2 is short and thick, and row 3 is short and thin. The short isoform of the molecular motor MYO15A (MYO15A-S) and the actin regulator EPS8 [6–8] couple with the scaffolding protein WHRN [9, 10] to form a complex with and transport the polarity proteins GPSM2 and GNAI3 [10, 11] to the tips of row 1 stereocilia, where all five proteins are stabilized. Formation of this complex controls elongation of the actin core, as bundles from genetic or functional knockouts of *Gpsm2, Gnai3, Eps8, Myo15a*, or *Whrn* have abnormally short stereocilia [10–14]. Row 2 is characterized by selective concentration of the heterodimeric capping protein subunit CAPZB, its partner TWF2, EPS8L2, and MYO15A-L, the long isoform of MYO15A [8, 15, 16].

Mechanotransduction is acquired by cochlear hair cells around the time of birth in mice and rats [17–19]. Transduction channels are relatively nonselective cation channels, with substantial permeation by Ca^2+^ [1]. The transmembrane channel-like proteins TMC1 and TMC2 make up at least part of the channel’s pore [20–22]. The small transmembrane protein TMIE, which may assist in transporting TMCs to stereocilia tips [23], is also absolutely required for transduction [24].

Several lines of evidence suggest that transduction regulates stereocilia dimensions. First, postnatal elimination of transduction after hair-bundle development leads to abnormally short row 2 and 3 stereocilia [25]. Moreover, blockade of transduction channels in cochlear cultures from early postnatal mice leads to a shortening of row 2 and 3 stereocilia over several hours [26]. Finally, while bundles of mice lacking expression of both TMC1 and TMC2 form relatively normally and have tip links [20], their morphology more closely resembles that of immature bundles [19].

Taken together, these results suggest that transduction may play a role in regulating development of the hair bundle. Here we show that the widening of row 2 and the narrowing of row 3 stereocilia both correlate with the acquisition of transduction, and that row 2 shortens and row 1 lengthens after the widening period has concluded. Correspondingly, stereocilia in mice lacking expression of both TMC1 and TMC2—or of TMIE—have nearly uniform row 1, 2, and 3 diameters and a smaller separation in height between all rows. Loss of transduction is also correlated with a loss of molecular row identity, so that MYO15A-S and MYO15A-L, as well as EPS8 and EPS8L2, are found at all stereocilia tips. At least for EPS8, this redistribution can be replicated with blockade of transduction channels in cochlear cultures. In addition, by the third postnatal week, GNAI3, GPSM2, CAPZB, and TWF2 failed to accumulate at tips, which could also contribute to the corresponding changes in stereocilia dimensions. We propose that like in chick cochlea, lengthening and widening steps are separated in time in mice, and that Ca^2+^ entry through transduction channels controls the diameter of rows 2 and 3 at a specific stage in bundle development and leads to coordination of stereocilia dimensions.

## Results

### Length and width of inner hair cell stereocilia during development

Because developmental changes in stereocilia dimensions have not been systematically investigated in the mammalian cochlea, we examined development in inner hair cells (IHCs) of C57BL/6J mice. We visualized stereocilia of apical IHCs for the following reasons: their large size allows accurate measurement of dimensions using fluorescence microscopy; their developmental window of interest is postnatal; dissection of older cochleas is easier at the apex; and the time course of transduction-current acquisition has been determined for apical cells [19]. Using phalloidin staining to visualize the actin cores of stereocilia (Fig. S1A), we generated x-z reslices of x-y-z stacks utilizing image-scanning microscopy with Airyscan detection (Fig. S1C), allowing us to examine a single column of stereocilia (rows 1, 2, and 3) in a hair bundle (Fig. 1A). We measured the apparent length and full width at half maximum (FWHM) of stereocilia during development (Fig. S1A); although the true dimensions are convolved with the microscope’s point-spread function (PSF), we achieved a lateral resolution of <200 nm (Fig. S1B). Representative examples of apical IHC stereocilia columns are shown in Fig. 1A for embryonic day 18.5 (E18.5) through postnatal day 19.5 (P19.5).

**Figure 1.**
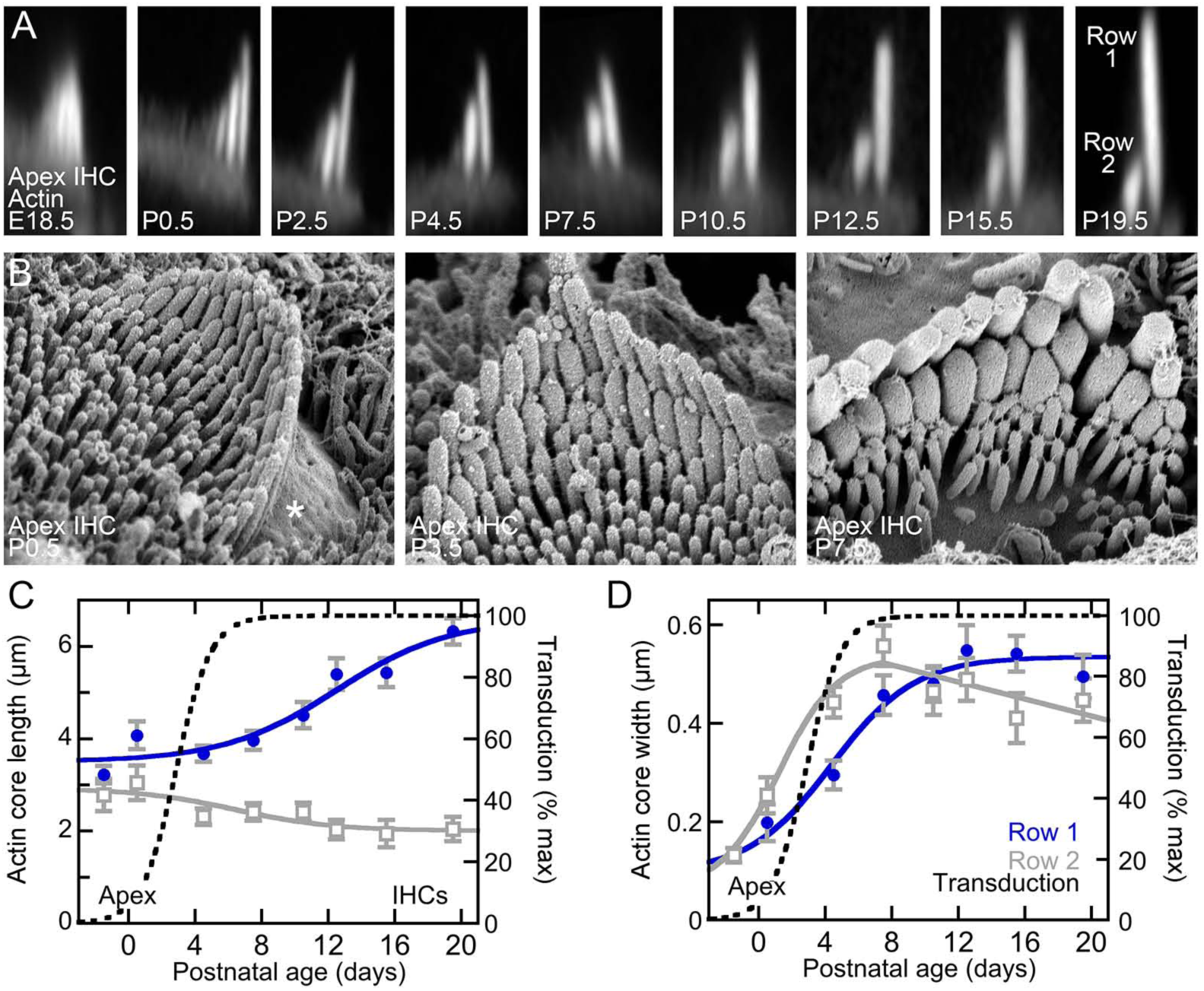
Length and width of apical inner hair cell stereocilia during development. ***A***, Examples of C57BL/6J stereocilia during development. Each panel is a x-z reslice taken from a phalloidin-stained IHC at the indicated age; all panels are 4 µm wide. ***B***, Scanning electron microscopy images of hair bundles from apical cochlea at P0.5, P3.5, and P7.5. Asterisk indicates the bare zone in the P0.5 panel. Panels are 5.2 µm wide. ***C***, Apparent actin core length for row 1 (blue) and row 2 (gray) during apical IHC development. Logistic-equation fit with midpoints of 12.5 (row 1) and 6.4 (row 2) days. ***D***, Apparent actin core width for rows 1 and 2 of apical IHCs during development. Logistic fit with midpoints of 4.4 (row 1) and 1.2 (row 2) days. Logistic fit to data from Beurg et al. (2018) for apical IHC transduction (midpoint of 2.9 days) was superimposed on both C and D. Data points indicate mean ± SD (n=21-99). See also Figure S1.

In contrast with previous reports [4, 5], we detected temporally distinguishable lengthening and widening stages that roughly corresponded to the phases of stereocilia growth originally reported for chick [3]. An initial lengthening step took place prior to E18.5, as row 1 stereocilia at the apex were already 3-4 µm tall by this time point (Fig. 1C). Rows 1 and 2 stereocilia did not change in length again until P4.5, when row 1 stereocilia began to lengthen and row 2 stereocilia shortened (Fig. 1C). Row 1 and row 2 stereocilia continued to lengthen and shorten, respectively, until their mature heights were reached by P21. As expected because of the basal-to-apical progression of hair cell development, row 1 lengthening and row 2 shortening occurred several days earlier in mid- and basal-cochlea regions (Fig. S1F, H).

As in chick cochlea hair cells, widening took place between the two lengthening phases of mouse IHCs (Fig. 1D). In apical hair cells, widening of row 2 of IHC stereocilia was correlated in time with acquisition of mechanotransduction (Fig. 1D). Row 2 widening also occurred several days earlier in mid- and basal-cochlea regions (Fig. S1G, I). Apical IHC row 1 widened as well, albeit delayed by several days, with the majority of widening occurring between P0.5 and P7.5 for apical IHCs (Fig. 1D). By P7.5, widened stereocilia in rows 1 and 2 were apparent by scanning electron microscopy (Fig. 1B). These data indicate that mouse IHC stereocilia develop with a pattern similar to that seen in chick cochlea [3]; widening of row 2 was correlated with transduction onset, and was followed by the simultaneous shortening of row 2 and lengthening of row 1 as the cells functionally matured.

### Morphology of hair bundles in mice lacking transduction

To test whether transduction affected stereocilia dimensions, we visualized hair bundles in cochleas from control mice and mice lacking one or both *Tmc* genes. In most experiments, we used *Tmc1*^*KO/KO*^;*Tmc2*^*KO*^/+ as controls, as one copy of the *Tmc2* gene was sufficient for hair-cell function and normal hair-bundle structure [19, 20]. As noted by Beurg and collaborators [19], IHC and OHC bundles in *Tmc1*^*KO/KO*^;*Tmc2*^*KO/KO*^ (*Tmc* DKO) mice had morphology that was altered as compared to controls (Fig. 2A-D). IHC stereocilia of *Tmc* DKO mice were arranged in a shallow ‘V’ shape (Fig. 2D). Three to five rows of similar-diameter stereocilia were present, and the staircase spacing between adjacent rows was considerably reduced compared to control bundles (Fig. 2K-L). OHC stereocilia of *Tmc* DKO mice were arranged in a ‘U’ shape, often with a disruption at the vertex of the ‘U’ (Fig. 2C); three to five rows were also present in OHCs, arranged in a shallow staircase.

**Figure 2.**
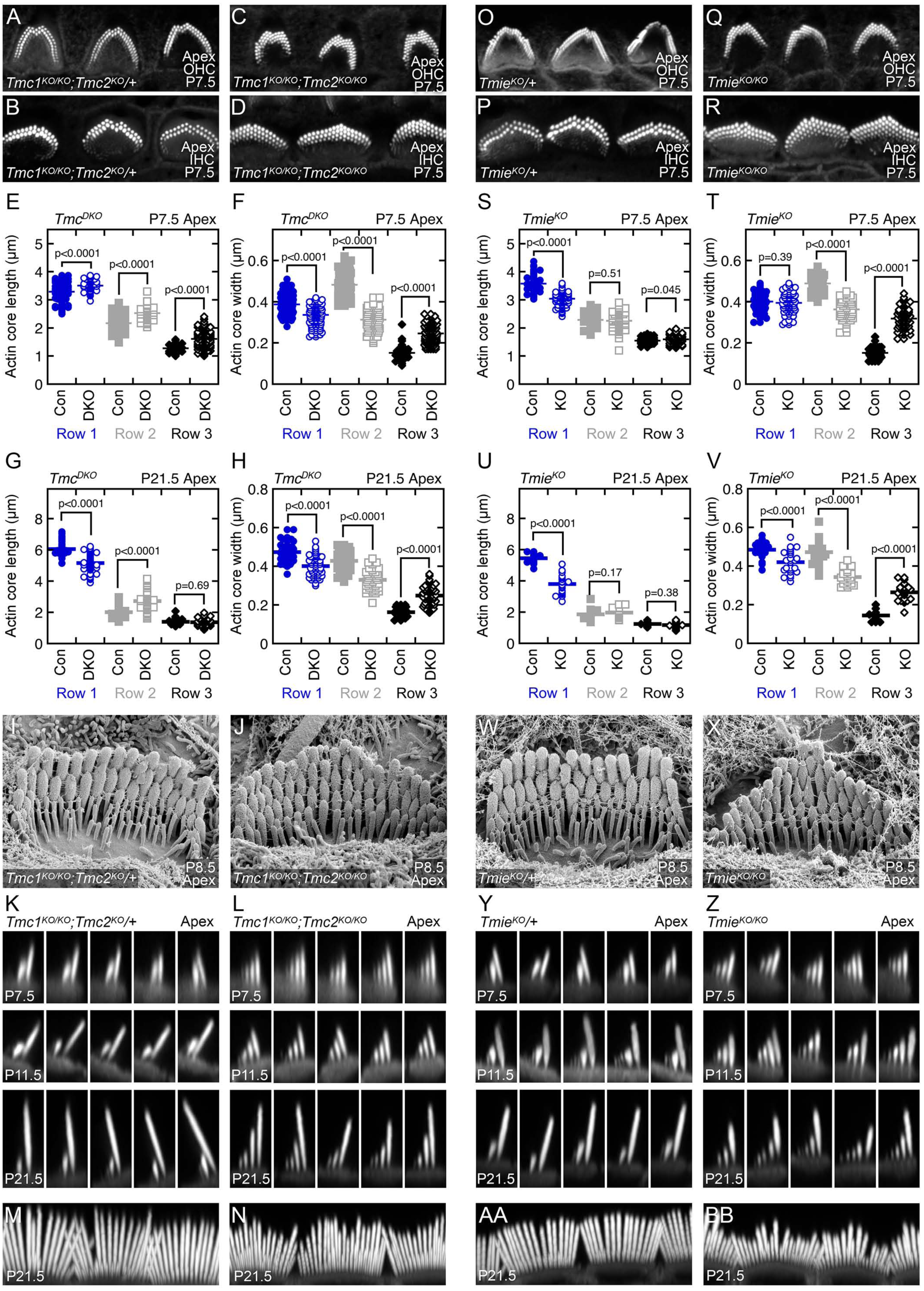
*Tmc* DKO and *Tmie* KO phenotypes. **A-N**, *Tmc* control and DKO. ***O-BB***, *Tmie* heterozygote and KO. ***A-D, O-R***, Phalloidin-stained images of apical hair bundles at P7.5 from *Tmc1*^*KO/KO*^;*Tmc2*^*KO*^/+ (A-B), *Tmc1*^*KO/KO*^;*Tmc2*^*KO/KO*^ (C-D), *Tmie*^*KO*^/+ (O-P), and *Tmie*^*KO/KO*^ (Q-R) outer and inner hair cells. Panels are 25 µm wide. ***E-H, S-V***, Stereocilia actin core length (E, G, S, U) and width (F, H, T, V) of *Tmc* (E-H) and *Tmie* (S-V) genotypes in apical bundles measured at P7.5 and P21.5. Pairwise comparisons used t-tests (P7.5: row 1, n=52-122; row 2, n= 52-122; row 3, n=36-122; P21.5: row 1, n=23-57; row 2, n=23-58; row 3, n=9-56). ***I-J, W-X***, Scanning electron microscopy images of *Tmc* (I-J) and *Tmie* (W-X) genotypes from apical cochlea at P8.5. Panels are 5.2 µm wide. ***K-L, Y-Z***, Reslice images (x-z) from bundles at the indicated ages from *Tmc* (K-L) and *Tmie* (Y-Z) genotypes. Each set shows different stereocilia ranks from a single hair bundle at that age. All panels are 4 µm wide. ***M-N, AA-BB***, Bundle en-face maximum projections of *Tmc* (M-N) and *Tmie* (AA-BB) genotypes from cochlea apex at P21.5. Panels are 25 µm wide. See also Figures S2 and S3.

We measured the length (Fig. 2E) and width (Fig. 2F) for IHC stereocilia of P7.5 control and *Tmc* DKO mice. While IHC rows 1 and 2 changed in length only modestly between control and *Tmc* DKO at this age, row 3 lengthened notably in mutants (Fig. 2E). Consistent with the correlation between transduction and row 2 widening (Fig. 1), row 2 was significantly less wide in stereocilia from *Tmc* DKO mice than from controls (Fig. 2F). Row 1 also narrowed but row 3 widened (Fig. 2F), which led to an overall equalization of *Tmc* DKO stereocilia widths.

Scanning electron microscopy of *Tmc* DKO hair bundles revealed the reorganized bundle structure and thickened row 3 and 4 stereocilia (Fig. 2I-J). The distribution of inter-stereocilia links, including those at positions suggesting that they were tip links, was similar between control and transduction mutant bundles (Fig. 2I-J).

While *Tmc2* is expressed early in cochlear hair-cell development, *Tmc1* only appears later [20]. Unlike *Tmc1*^*KO/KO*^;*Tmc2*^*KO*^/+ mice used as controls, mice that were homozygous for the *Tmc2* knockout but heterozygous for the *Tmc1* knockout (*Tmc1*^*KO*^/+;*Tmc2*^*KO/KO*^) had a hair-bundle phenotype that was intermediate between wild-type and *Tmc* DKO bundles (Fig. S2A-D). Many IHC bundles appeared normal, but others had a few thicker row 3 stereocilia, which were usually found in the middle of the row. Other bundles had complete rows of thick row 3 stereocilia (Fig. S2D). The lack of TMC molecules during early development in *Tmc1*^*KO*^/+;*Tmc2*^*KO/KO*^ mice thus leads to a partial morphology phenotype.

*Tmie*^*KO/KO*^ (*Tmie* KO) largely phenocopied *Tmc1*^*KO/KO*^;*Tmc2*^*KO/KO*^ stereocilia dimensions and arrangement at P7.5 (Fig. 2O-R). Lengths of P7.5 *Tmie* KO stereocilia rows were similar to those of *Tmc* DKOs (Fig. 2S), as were widths (Fig. 2T). Scanning electron microscopy revealed not only the altered stereocilia length and width (Fig. 2W-X), but also elongated stereocilia tips, as previously reported [24].

We also compared *Tmc* and *Tmie* mutants later in development (Fig. 2G-H, K-N, U-V, Y-BB). Differences in diameter between control and mutant stereocilia remained at P21.5 (Fig. 2H, V). As cochleas matured, both transduction mutants showed an increasing variability of the lengths of row 1 stereocilia, especially obvious at P21.5 (Fig. 2G, N, U, BB). There were significant differences between the two mutants, however. Heights of *Tmc* DKO row 1 stereocilia were on average close to control heights, although there was some length variability in rows 1 and 2 as the hair bundle matured (Fig. 2G, N). By contrast, row 1 stereocilia of *Tmie* KO IHCs were shorter and thicker than control row 1 stereocilia, particularly obvious at P21.5 (Fig. 2U, BB); often 3-4 stereocilia of a bundle were much thicker than all others in a bundle (Fig. 2BB). In addition, *Tmie* KO row 1 stereocilia were significantly more variable in their length (Fig. 2U, BB). Row 1 stereocilia of *Tmie* controls at P21.5 were 5.47 ± 0.29 µm long (mean ± SD; n=23), for a coefficient of variation of 0.05. By contrast, the coefficient of variation for *Tmie* KO row 1 stereocilia was 0.16, more than three-fold greater (3.77 ± 0.58 µm; n=23). As length measurements were performed mostly on central stereocilia, this variability may be an underestimate. Together, these results indicate that transduction is necessary to create the steep staircase architecture with distinct row widths that is characteristic of IHC bundles, as well as to maintain coordinated stereocilia dimensions across each row as the cells mature. Because of the phenotypic distinction between *Tmc* DKO and *Tmie* KO bundles, TMIE may play an additional role in bundle development beyond the targeting of the TMCs.

We also examined stereocilia dimensions in *Myo15a*^*sh2*^ mice, which have a mutation that inactivates the motor’s ATPase; abundant short stereocilia typify this mutant [12, 27]. At P7.5, lengths and widths of *Myo15a*^*sh2/sh2*^ stereocilia of inner hair cells were very similar across all rows (Fig. S3A-B). Scanning electron microscopy showed the altered length, width, and number of stereocilia in the *sh2/sh2* mutants, as well as an abundance of interstereocilia links (Fig. S3C-D).

### Developmental appearance of row 1 identity proteins

We next examined whether the changes in stereocilia dimensions in mice lacking transduction were correlated with changes in the localization of protein complexes specific for row 1 or 2, which can regulate the actin cytoskeleton. In wild-type cochlear hair cells, row 1 identity and stereocilia length are set by GNAI3- and GPSM2-dependent localization of a complex containing EPS8, MYO15A-S, and WHRN [10, 11, 14]. We localized each protein by immunofluorescence (Fig. 3A) and also quantified the fluorescence signal at individual IHC row 1 and row 2 tips (Fig. 3B). For four developmental ages—P0.5, P7.5, P15.5, and P21.5—we reported the average fluorescence at the tips of rows 1 and 2 combined, normalized to the peak average tip fluorescence, labeled “Avg %”; this measure (top panels) gives an approximate indication of total expression level at stereocilia tips. We also reported the fluorescence for each row 1 or row 2 tip divided by the average for that hair bundle, labeled “Tip / avg,” which reports the relative concentration of each protein at row 1 or 2 tips.

**Figure 3.**
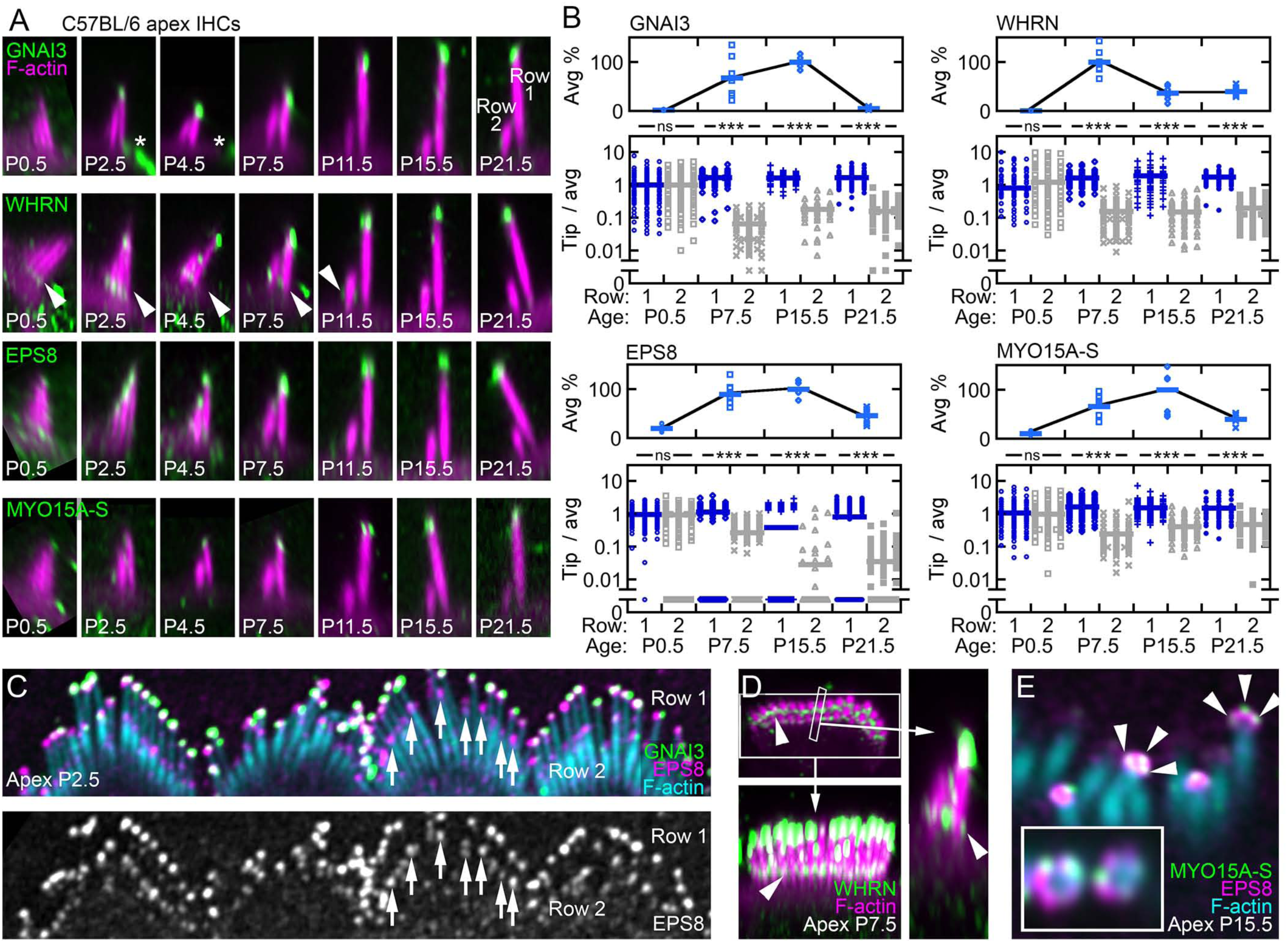
Distribution of row 1 proteins during development of C57BL/6J stereocilia. ***A***, Localization of proteins specific for row 1 stereocilia tips in IHC hair bundles during postnatal development. All images are x-z reslices. Brightness for antibody signal (green) and phalloidin for actin (magenta) was adjusted to represent the range in the image; intensities should not be compared between panels, even for the same protein. Similar panels with fixed antibody brightness are in Fig. S4B. All panels are 4 x 7.5 µm; in some cases, images were expanded with black background to fill the panel. Bare-zone labeling for GNAI3 is indicated by asterisks; arrowheads indicate ankle-link labeling for WHRN. ***B***, Quantitation of row 1 proteins in IHC bundles during development. Top panels: average signal for row 1 and 2 tips combined. Immunoreactivity for all row 1 and 2 tips for each bundle were plotted as percentage of the peak time point (n=4-9 per time point); solid bars indicate average for each time point. Bottom panels: ratio of immunoreactivity in individual row 1 or row 2 tips to the average immunoreactivity for both rows; solid bars indicate average. ****, p<0.0001; ns, not significant. Number of row 1 or row 2 stereocilia counted for each condition (n for bottom panels): GNAI3, 48-146; EPS8, 36-183; MYO15A-S, 53-138; WHRN, 65-134. ***C***, EPS8 labeling at row 2 tips during early development (P2.5). Arrows indicate examples of EPS8 at row 2 tips in one bundle. ***D***, Whirlin ankle-link labeling (arrowheads) at P7.5. Upper left image is horizontal slice (Fig. S1E) at the level of the ankle links; panel is 9 µm wide. Lower left panel is en face view (Fig. S1D) showing stereocilia; panel is 9 µm wide. Right panel is x-z reslice (small box); panel is 2.5 µm wide. ***E***, Antibody and phalloidin labeling shows the distribution of MYO15A-S (arrowheads) and EPS8 at stereocilia tips. MYO15A-S punctae are smaller than the EPS8 blob. See also Figure S4.

GNAI3 was not present at IHC stereocilia tips at P0.5, but began to accumulate at row 1 tips as early as P2.5 (Fig. 3A). For Fig. 3A, acquisition settings were kept constant but brightness settings were adjusted to reveal staining patterns at each age; examples of row 1 proteins over development with identical brightness settings are displayed in Fig. S4B. GNAI3 total signal increased at row 1 tips to a peak at P15.5, then dropped to low levels by P21.5 (Fig. 3B). GPSM2 showed a nearly identical labeling pattern with GNAI3 (Fig. S5A-B). WHRN showed a similar pattern of expression, with protein only starting to accumulate at row 1 tips at P2.5; WHRN peaked at row 1 tips during the second postnatal week, then declined by P21.5 (Figs. 3A-B, S4B). As shown previously [10, 28, 29], WHRN was also found in the ankle link region (Fig. 3A,D), while GNAI3 and GPSM2 were also found within the apical membrane’s bare zone during the first postnatal week (Fig. 3A).

EPS8 also increased at IHC stereocilia tips between P0.5 and P15.5, then declined by P21.5 (Figs. 3A-B, S4B). Unlike GNAI3 and WHRN, EPS8 was present at stereocilia tips at P0.5 and was at substantial levels at both row 1 and row 2 tips at early ages (Fig. 3C), only becoming fully restricted to row 1 after P7.5. As previously demonstrated [10], the expression pattern for MYO15A-S closely resembled that of EPS8 (Figs. 3A-B, S4B). We localized MYO15A-S with the TF1 antibody [30]. Although TF1 binds an epitope shared by MYO15A-S and MYO15A-L (Fig. S4A), comparison of its staining pattern to those of PB48, a pan-MYO15A antibody [30], and PB888, an antibody specific for MYO15A-L [8, 30], indicated that TF1 is likely specific for MYO15A-S (Fig. S4C). Like EPS8, MYO15A-S was present at IHC stereocilia tips in P0.5 apical hair cells, with labeling at both row 1 and row 2 tips until P4.5 that became increasingly restricted to row 1 between P7.5 and P21.5 (Fig 3A,B). MYO15A-S and EPS8 had a similar expression pattern, with increased tip accumulation between P0.5 and P15.5 that declined by P21.5 (Fig 3B).

In contrast to previous reports [10], we found in some of our images that EPS8 and MYO15A-S did not have perfectly overlapping distribution at row 1 tips. EPS8 usually formed a cap that encompassed the entire tip and extended down a few hundred nanometers (Fig. 3A, E). By contrast, MYO15A-S was sometimes present in 1-3 bright puncta, each of which was smaller than the EPS8 cap (Fig. 3E).

Appearance of GNAI3 and WHRN at row 1 stereocilia tips, starting at P2.5, is thus correlated with the onset of transduction. EPS8 and MYO15A-S localize to stereocilia tips prior to transduction onset, however, and become fully restricted to row 1 only after transduction onset. Each of these proteins peak in their selective accumulation at row 1 tips between P7.5 and P15.5, when row 1 undergoes the majority of its lengthening, and have declined by 21.5, when the stereocilia have reached their mature dimensions (Figs. 3A-B, S4B).

### Loss of row 1 identity in mice lacking transduction

In *Tmc* DKO or *Tmie* KO mice, proteins specific for row 1 had substantially altered distributions (Fig. 4). Both EPS8 and MYO15A-S redistributed from row 1 only in controls to all rows in *Tmc* DKO and *Tmie* KO IHCs, albeit more strongly in *Tmie* KO hair cells (Fig. 4). The redistribution was relatively modest at P7.5 (Fig. 4C-D, G-H), with scattered appearance of each protein at row 2 tips in the mutants, but was robust by P21.5 (Fig. 4K-L, O-P).

**Figure 4.**
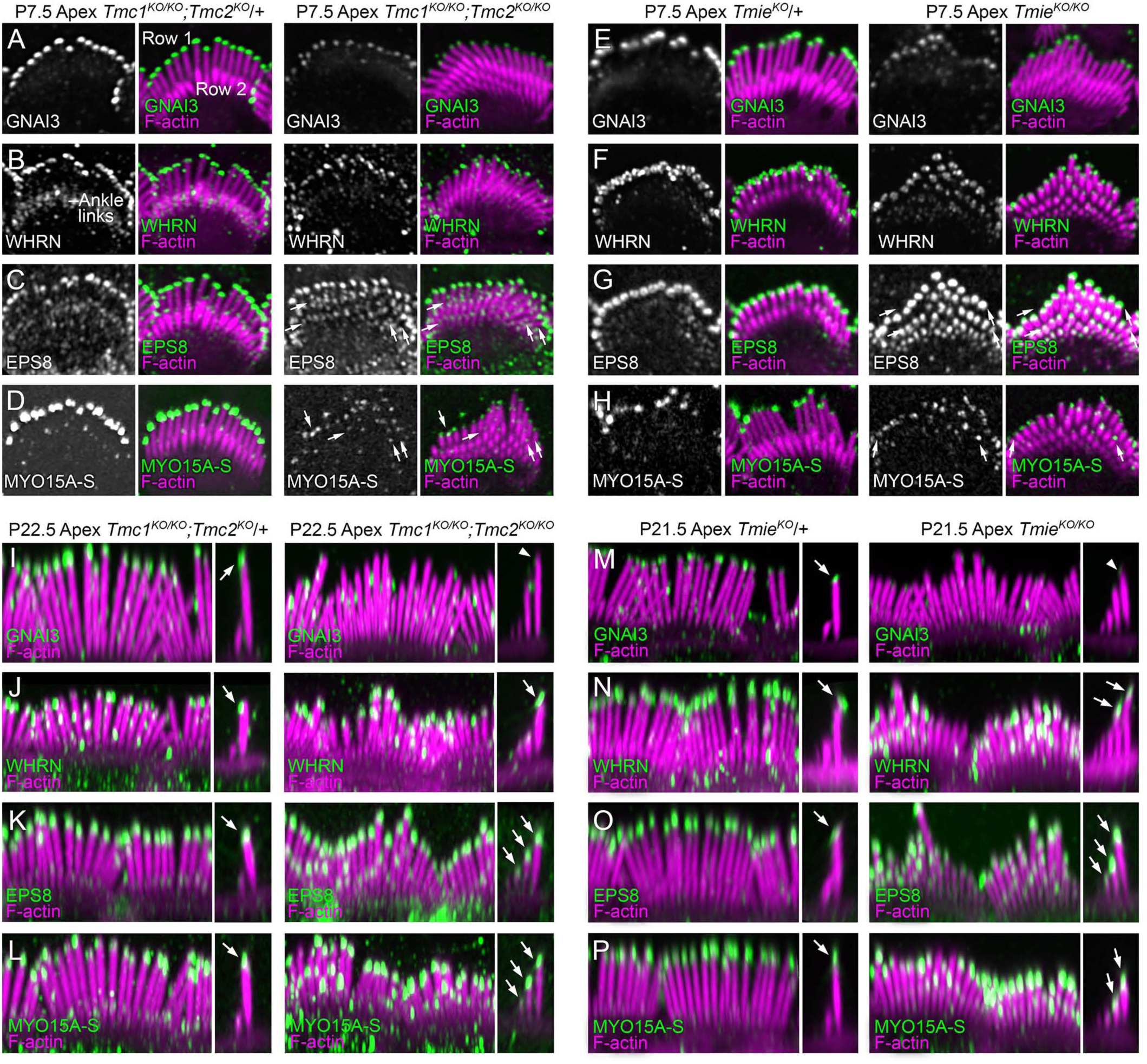
Redistribution of row 1 proteins in transduction mutants. ***A-H***, Row 1 tip proteins at P7.5 in *Tmc* DKO and *Tmie* KO mice. Left paired images are antibody channel and antibody-actin merge for control IHCs; right paired images are for DKO or KO mice. WHRN ankle-link labeling is indicated in D. Arrows indicate examples of labeling at stereocilia tips of rows 1-3 in mutant hair bundles. Panels are 7.5 µm wide. ***I-P***, Row 1 tip proteins at P22.5 in *Tmc* DKO and P21.5 in *Tmie* KO mice. En face views, 15 µm wide (left), and x-z reslices, 4 µm wide (same scale; right). Arrowheads in I and M indicate the decrease seen in GNAI3 by P21.5; arrows in panels I-P indicate staining at tips of row 1 (in control IHCs) or tips of rows 1-3 (in transduction mutants). Each pair of control and DKO or KO panels was from littermates; during image acquisition, the gain setting for the antibody signal was adjusted to reveal the staining pattern in each control sample, and the corresponding DKO or KO sample used the same gain setting. For display, brightness settings for the panels were kept constant between control and mutant. See also Figure S5.

We quantified row 1 protein distribution at the tips of rows 1 and 2 in hair bundles of control and transduction-mutant IHCs (Fig. 5). Behavior of the four row 1 proteins was similar between *Tmc* DKO and *Tmie* KO IHCs. At P7.5, average tip fluorescence levels decreased significantly for GNAI3 and WHRN (Fig. 5A, C, I, K), but were unchanged for EPS8 and MYO15A-S (Fig. 5E, G, M, O). For EPS8 and MYO15A-S, the normalized row 2 signal increased ∼2-fold in *Tmc* DKO or *Tmie* KO IHCs compared to controls, with the normalized row 1 signal decreasing accordingly (Fig. 5F, H, N, P). While the GNAI3 normalized signal at row 2 also increased (Fig. 5B, J), this may not represent redistribution as the GNAI3 signal level at row 2 was close to background levels. The normalized row 2 signal for WHRN did not change between controls and mutant IHCs (Fig. 5D, L). The *Tmc1*^*KO/KO*^;*Tmc2*^*KO*^/+ IHCs we used as controls had a slight redistribution phenotype for EPS8 and decrease in row 1 GNAI3 localization at P7.5, similar to *Tmc1*^*KO*^/+;*Tmc2*^*KO/KO*^ cochleas, but redistribution and signal reduction (for GNAI3) was stronger in *Tmc* DKO cochleas (Fig. S5K-R).

**Figure 5.**
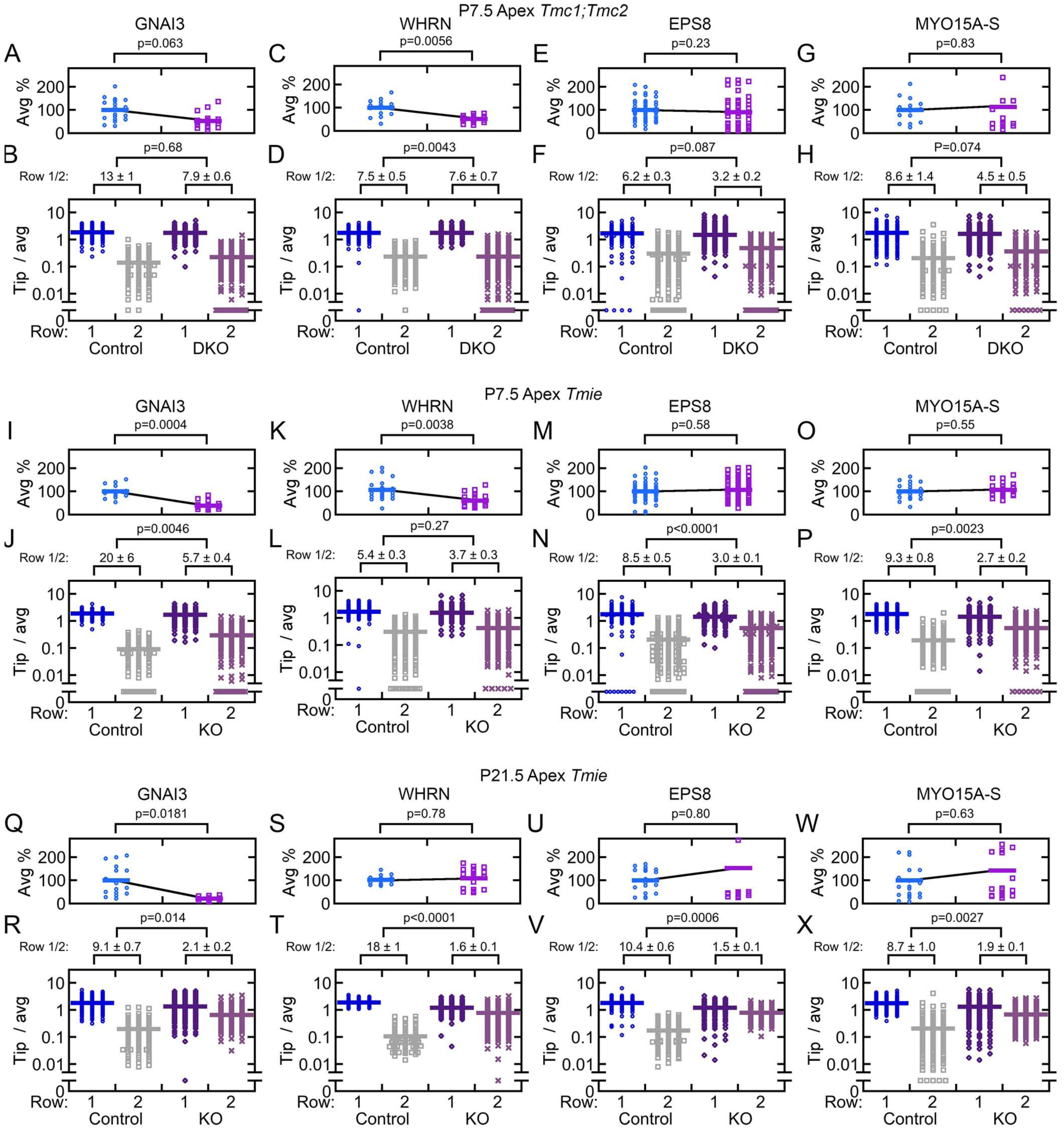
Quantitation of redistribution of row 1 proteins in transduction mutants. Quantitation was carried out as described for Fig. 3. For each protein for a given genotype, the upper panel shows the average fluorescence for row 1 and 2 tips from each hair bundle, normalized to average control fluorescence for the experiment. Pairwise control vs. mutant comparisons used the Student’s t-test (p values indicated above panel). The bottom panels show row 1 or row 2 fluorescence normalized to the average row 1 + 2 fluorescence for each bundle. The average ratio of row 1 tip/average to row 2 tip/average (mean ± SEM) is indicated for each condition. ***A-H***, P7.5 *Tmc* control and DKO distribution of GNAI3 (A-B), EPS8 (C-D), MYO15A-S (E-F), and WHRN (G-H). ***I-P***, P7.5 *Tmie* control and KO distribution of GNAI3 (I-J), EPS8 (K-L), MYO15A-S (M-N), and WHRN (O-P). ***Q-T***, P21.5 *Tmie* control and KO distribution of GNAI3 (Q-R), EPS8 (S-T), MYO15A-S (U-V), and WHRN (W-X). Bundles quantified for each condition (n): A, 19-20; C, 50-55; E, 15; G, 16-20; I, 16-20; K, 62-68; M, 23-24; O, 23-24; Q, 16; S, 19; U, 20; W, 18. Row 1 or row 2 stereocilia counted for each condition (n): B, 190-200; D, 338; F, 150; H, 160-200; J, 160-200; L, 408-457; N, 230-240; P, 240; R, 192; T, 228; V, 240; X, 216.

When quantified at P21.5, the redistribution was even more clear (Fig. 5Q-X). The normalized GNAI3 and WHRN fluorescence at row 2 tips increased more than four-fold between *Tmie* control and KO stereocilia (Fig. 5R, T). Likewise, normalized EPS8 and MYO15A-S signal at row 2 tips increased ∼five-fold between control and KO stereocilia (Fig. 5V, X). Total intensity at rows 1 and 2 was reduced substantially in *Tmie* KO IHCs for GNAI3 (Fig. 5Q), but was unchanged for WHRN, EPS8, and MYO15A-S (Fig. 5S, U, W).

*Tmc1*^*KO/KO*^;*Tmc2*^*KO*^/+ mice become deaf after TMC2 disappears [20]. At P21.5, a partial phenotype was apparent in *Tmc1*^*KO/KO*^;*Tmc2*^*KO*^/+ IHCs, which should have no transduction at this age; the redistribution of GNAI3 and EPS8 was intermediate between that of the *Tmc1*^*KO*^/+;*Tmc2*^*KO*^/+ double heterozygotes and the *Tmc1*^*KO/KO*^/+;*Tmc2*^*KO.KO*^ double knockouts (Fig. S5S-V). Although they may be involved in stereocilia lengthening and widening and have to been shown to be present at all stereocilia tips [31, 32], there was no obvious change in tip localization of MYO3A, ESPN-1, and ESPNL in *Tmc* DKO IHCs (Fig. S5E-J).

Together, these results all suggest that as row 1 lengthens between P7.5 and P21.5, transduction promotes accumulation of GNAI3, WHRN, EPS8, and MYO15A-S at row 1 tips. In transduction mutants, redistribution of row 1 proteins to all stereocilia rows and the loss of GNAI3 accumulation at row 1 tips could both contribute to reduced row 1 lengthening and row 2 shortening.

### Redistribution of row 1 proteins in the absence of Ca^2+^ entry

We hypothesized the normal lack of EPS8 and MYO15A-S at row 2 and 3 tips in wild-type IHCs could be because Ca^2+^ entry through active transduction channels prevents their accumulation there. If so, blockade of transduction, which prevents Ca^2+^ entry, should cause their redistribution. Using an established paradigm [26], we cultured P4.5 C57BL/6J cochleas for 24 hr in vitro with a control solution or the same solution containing 100 µM tubocurarine, an inhibitor of transduction [33]. Under control conditions, FM1-43, a permeant channel blocker and fluorescent marker [34, 35], labeled cochlear hair cells, albeit OHCs more strongly than IHCs (Fig. 6A). In the presence of tubocurarine, however, labeling of both types of hair cells by FM 1-43 was substantially reduced, confirming the effective blockade of transduction channels (Fig. 6B).

**Figure 6.**
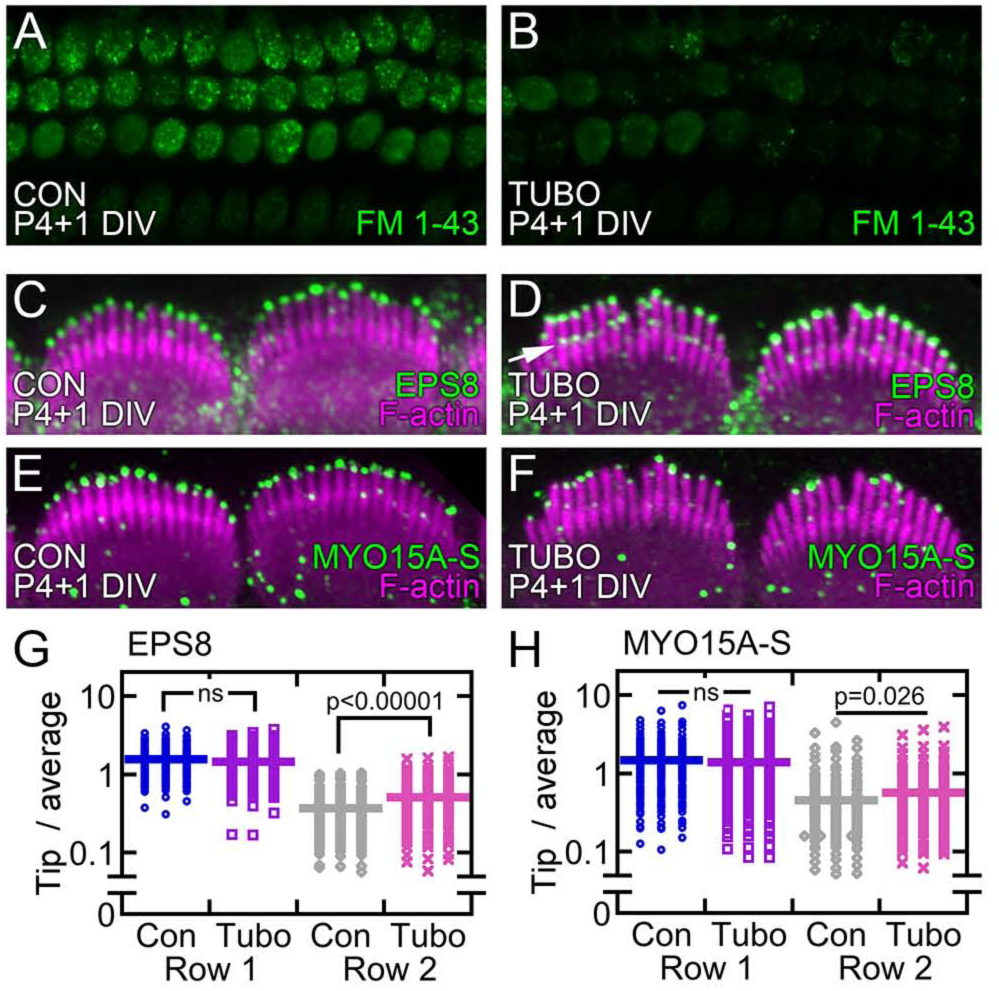
Blockade of transduction channel causes partial redistribution of EPS8 and MYO15A-S. ***A-B***, FM1-43 assessment of transduction in cochlear cultures treated for 24 hr with control (A) and tubocurarine (B) solutions. Dye loading of OHCs and IHCs was diminished by tubocurarine (TUBO). Panel full widths of 50 µm. ***C-D***, EPS8 immunoreactivity in cochlear cultures treated for 24 hr with control (A) and tubocurarine (B) solutions. EPS8 labeling increased at row 2 stereocilia tips (white arrow). ***E-F***, MYO15A-S immunoreactivity in control (E) and tubocurarine-blocked (F) cochlear cultures. Panels C-F are 18 µm wide. ***G-H***, Quantitation of EPS8 (G) and MYO15A-S (H) redistribution. Vertical axis represents ratio of immunoreactivity in row 1 or row 2 to the average immunoreactivity in both rows; the mean is represented by solid bar. Significance by t-test is indicated above.

When we stained cochlear cultures with antibodies against row 1 proteins, we noted a partial redistribution to row 2 (Fig. 6C-F). There was considerable variability among hair bundles, however, and relocalization to row 2 was more apparent with EPS8 (Fig. 6D) than with MYO15A-S (Fig. 6F). Quantitation of fluorescence intensity at row 1 and row 2 tip showed that tubocurarine significantly increased the EPS8 staining at row 2 (Fig. 6G). MYO15A-S also was present at higher levels on row 2 tips, albeit with more variability that lead to a considerably higher p value (Fig. 6H). These experiments suggest that the effect of transduction on row 1 protein distribution is through ions passing through the channel, presumably Ca^2+^, and not due to the presence of the transduction complex proteins themselves.

### Developmental appearance of row 2 identity proteins

We next examined the time course of accumulation of proteins specific to tips of stereocilia row 2. As previously reported [15], CAPZB appeared along apical IHC shafts at early ages, with little distinction between rows 1 and 2 (Fig. 7A-B); at P7.5, CAPZB often appeared to be associated with the membrane or periphery of the actin core (Fig. 7E), although that pattern of localization that could be due to poor antibody penetration [36]. At P15.5 or later, CAPZB was strongly concentrated at row 2 tips (Fig. 7A, arrow; Fig. 7F, arrowhead). TWF2 showed a similar pattern to that of CAPZB at P15.5 and later, but was located at ankle links between P0.5 and P7.5 (Fig. 7A, arrowheads). Using *Twf2*^*KO/KO*^ mice, we verified that the TWF2 staining in the ankle-link region was specific (Fig. S6A-B). At P15.5, while observed at some row 2 tips (Fig. 7D, arrowheads), TWF2 was also seen in row 1 stereocilia shafts (arrows).

**Figure 7.**
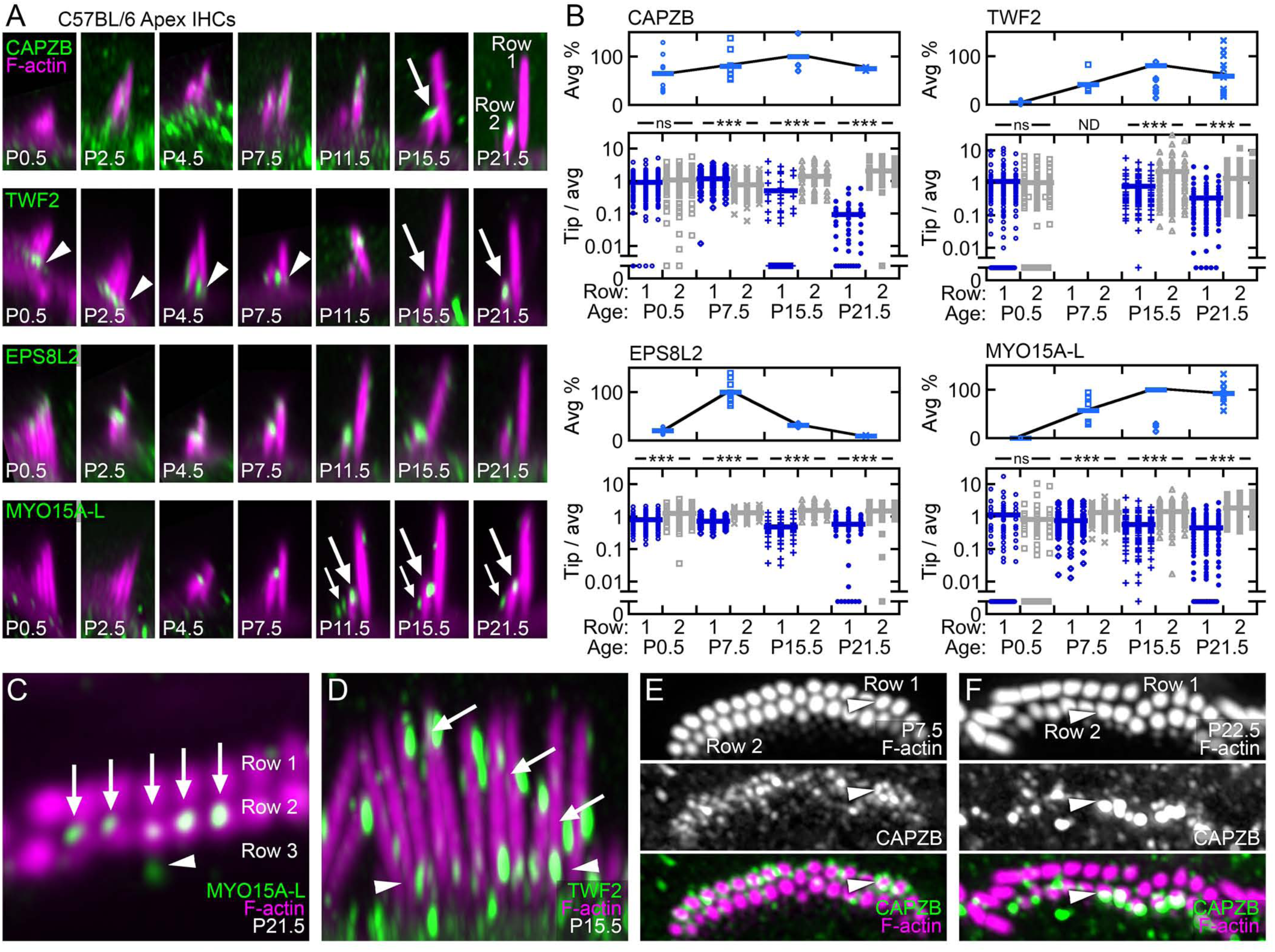
Distribution of row 2 proteins during development of C57BL/6J stereocilia. ***A***, Localization of proteins specific for row 2 stereocilia tips in IHC bundles during postnatal development. Brightness settings for the antibody signal (green) and phalloidin for actin (magenta) were adjusted for each panel to represent the range in the image; intensities should not be compared between panels. All panels are 4 x 7.5 µm; in some cases, images were expanded with black background to fill the panel. ***B***, Quantitation of row 2 proteins in IHC bundles during development. Quantitation was carried out as in Fig. 3. TWF2 was not measured at P7.5 (ND) as ankle-link and tip staining often overlapped in x-y sections. Number of row 1 or row 2 stereocilia counted for each condition (n): CAPZB, 43-134; EPS8L2, 47-160; MYO15A-L, 57-130; TWF2, 83-124. ***C***, MYO15A-L (arrows) at row 2 stereocilia tips. One row 3 tip was also captured (arrowhead). MYO15A-L punctae are located on the side of the row 2 tip where the tip link base would be located. ***D***, TWF2 at P15.5, between early ages when TWF2 is at ankle links and late ages when TWF2 is at row 2 tips. Arrows indicate strong labeling in row 1 stereocilia shafts; arrowheads indicate labeling at row 2 tips. ***E-F***, CAPZB at P7.5 (E) and P22.5 (F). Arrowheads indicate labeling near row 1 membranes in E and at row 2 tips in F. Panel full widths: C, 5 µm; D, 9.1 µm; E-F, 10 µm. See also Figure S6.

Unlike CAPZB and TWF2, EPS8L2 was clearly seen at IHC stereocilia tips at P0.5, and became concentrated at row 2 tips as early as P2.5 (Fig. 7A). The EPS8L2 row 2 to row 1 enrichment became more pronounced after P7.5, but was never large; a low EPS8L2 signal could be seen at row 1 stereocilia tips at all ages (Fig. 7A, B). Immunocytochemistry for EPS8L2 was variable, however, with most preparations showing concentration at tips but occasional samples showing broader stereocilia distribution. At P15.5 and later, CAPZB, TWF2, and EPS8L2 were apparent at row 2 but not row 3 tips.

As described previously [8], MYO15A-L did not appear in IHC stereocilia until P4.5 (Fig. 7A,B). MYO15A-L was seen in large punctae at row 2 tips, often on the side of row 2 stereocilia tips where a tip link should connect to the taller stereocilum (large arrows in Fig. 7A, C). MYO15A-L was also strongly detected at the tips of the very short row 3 stereocilia (Fig. 7A, small arrows). The pan-MYO15A PB48 antibody detected MYO15A at tips of all rows of stereocilia, as expected (Fig. S4C). Compared to row 1 specific proteins, proteins specific for row 2 accumulate later on in development with the mature pattern only appearing after P15.5.

### Loss of row 2 identity in mice lacking transduction

Distribution of row 2 proteins was also altered in mutant mice lacking transduction. In *Tmie* KOs at P21.5, rather than being concentrated at row 2 tips as in controls, CAPZB was absent from all tips and was found along the shafts of all stereocilia rows, maintaining a similar distribution to that seen earlier in development (Fig. 8A, E). By contrast, TWF2 was largely absent from P21.5 *Tmie* KO stereocilia (Fig. 8B, F). EPS8L2 shifted from concentration only at row 2 to significant presence at row 3 and even row 4 as well, albeit still little at row 1 tips (Fig. 8C), matching the distribution of MYO15A-L (Fig. 8D). Both EPS8L2 and MYO15A-L were found along row 1 shafts at higher levels in *Tmie* KOs as compared to controls (Fig. 8C, D).

**Figure 8.**
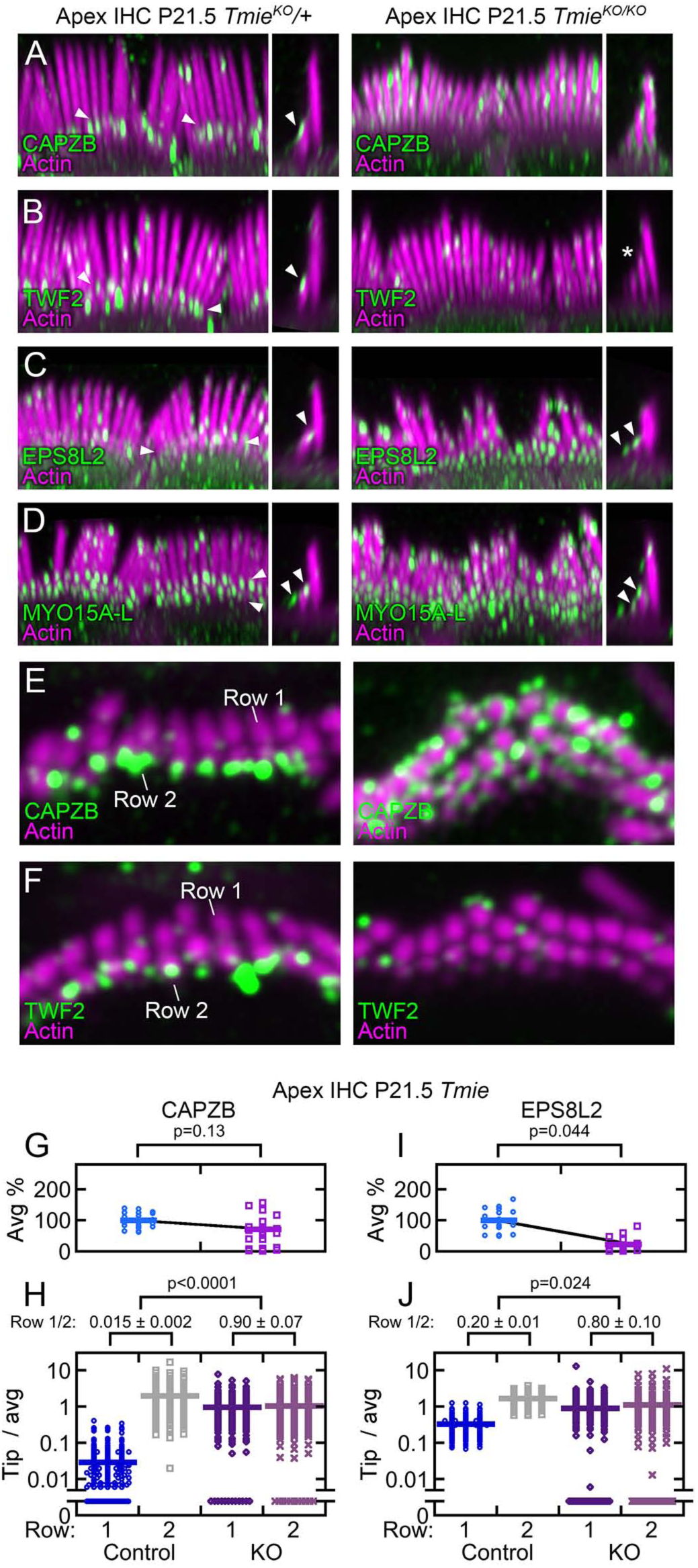
Redistribution of row 2 proteins in transduction mutants. ***A-B***, Views of row 2 proteins in *Tmie* control (left) and *Tmie* KO (right) samples. CAPZB (A), EPS8L2 (B), MYO15A-L (C), and TWF2 (D). Each pair of control and KO panels were from littermates, with identical sample preparation and confocal imaging parameters. Gain during acquisition and brightness of displayed panels were maintained as described earlier. Left panels are en face views, 15 µm wide (left), and right panels are x-z reslices, 4 µm wide (same scale). Arrowheads indicate staining at row 2 tips and, in some cases, at row 3 tips. The asterisk in the TWF2 panel for *Tmie* KO indicates that signal is reduced compared to control. ***E-F***, Horizontal slices at the level of row 2 tips for CAPZB (E) and TWF2 (F) in P21.5 *Tmie* KO. CAPZB labeling surrounds stereocilia shafts in mutants (E), while TWF2 labeling is absent in mutants (F). Panels are 7 µm wide. ***G-J***, Quantitation of CAPZB (G-H) and EPS8L2 (I-J) in P21.5 *Tmie* KO. Quantitation was performed as in Fig. 3. See also Figure S7.

*Tmc1*^*KO/KO*^;*Tmc2*^*KO*^/+ mice had a partial row 2 redistribution phenotype by P21.5. For example, CAPZB labeling was altered in *Tmc1*^*KO/KO*^;*Tmc2*^*KO*^/+ stereocilia; in this genotype, the redistribution of CAPZB to row 1 was as substantial as it was in the *Tmc1*^*KO/KO*^;*Tmc2*^*KO.KO*^ double knockouts (Fig. S7E-F). Similarly, EPS8L2 labeling was strong at row 2 and 3 tips of P21.5 *Tmc* DKO IHCs (Fig. S7G-H).

MYO15A could mediate some of the effects of transduction, e.g., through Ca^2+^ binding. Although EPS8L2 was reported to be at tips of vestibular stereocilia in *Myo15a*^*sh2/sh2*^ mice [16], its localization in IHCs was more complex. At P7.5, EPS8L2 was robustly expressed in *Myo15a*^*sh2/sh2*^ stereocilia, but was found along the shafts and was not at tips (Fig. 9A). By P15.5, however, EPS8L2 was largely absent from *Myo15a*^*sh2/sh2*^ IHC stereocilia (Fig. 9B). By contrast, CAPZB and TWF2 were seen prematurely at all *Myo15a*^*sh2/sh2*^ stereocilia tips, at P7.5 (Fig. 9C), P15.5 (Fig. 9D), or P21.5 (Fig. 9E-F).

**Figure 9.**
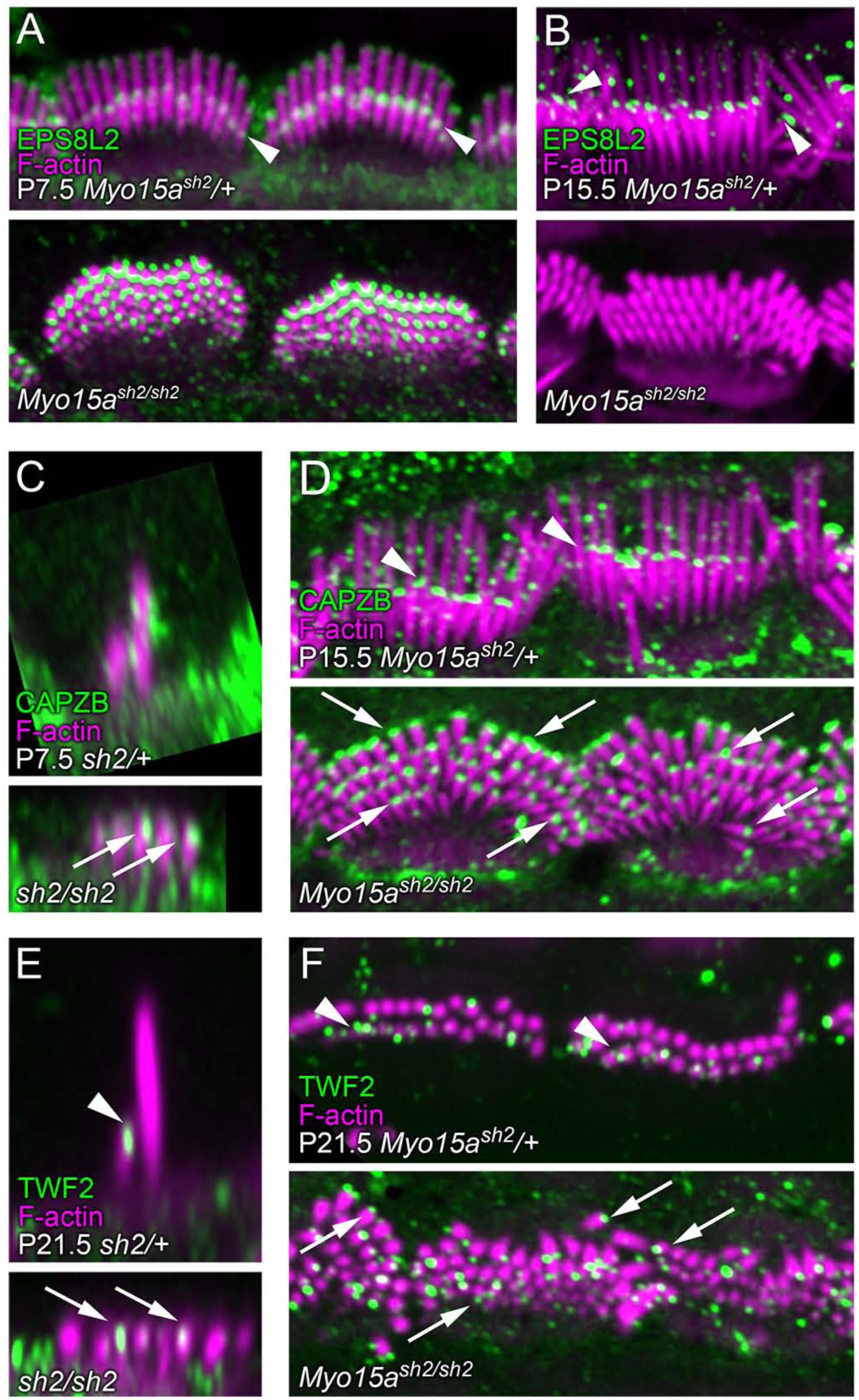
Redistribution of actin-capping proteins in *Myo15a*^*sh2/sh2*^ mice. Immunolocalization in *Myo15a*^*sh2*^*/+* and *Myo15a*^*sh2/sh2*^ apical IHC hair bundles. Arrows indicate row 1 labeling and arrowheads indicate row 2 labeling. Each pair of control and KO panels were from littermates, with identical sample preparation and confocal imaging parameters. Gain during acquisition and brightness of displayed panels were maintained as described earlier. ***A***, EPS8L2 at P7.5. ***B***, EPS8L2 at P15.5. ***C***, CAPZB at P7.5. ***D***, CAPZB at P15.5. ***E***, TWF2 at P7.5. ***F***, TWF2 at P21.5. Panels are 7 µm (G-I), 12.5 µm (K), or 20 µm (E-F, J) wide. Images are x-z reslice (G-I), flat (E, J-K), or horizontal (F) views. In some panels, *Myo15a*^*sh2*^*/+* and *Myo15a*^*sh2/sh2*^ are abbreviated *sh2/+* and *sh2/sh2*.

Taken together, analysis of row 2 proteins in *Tmc* and *Tmie* mutants shows that transduction not only upsets the molecular identity of row 1, but also leads to the redistribution of row 2 proteins. Transduction is thus central to defining the molecular polarity of the hair bundle.

## Discussion

Tilney suggested that Ca^2+^ entry through mechanotransduction channels regulates elongation of stereocilia [37]. We find that during development, the presence of key molecules underlying transduction are necessary for both normal lengths and widths of the shorter (transducing) rows of stereocilia. Specifically, in hair bundles of inner hair cells, the widening of row 2 and the thinning of row 3 were correlated temporally with the developmental appearance of transduction. Consistent with those results, in transduction-lacking *Tmc* DKO or *Tmie* KO IHCs, stereocilia lengths and widths were more uniform between the three rows. Moreover, the molecular identity of IHC rows—as assessed by tip-protein distribution in mutant bundles—was substantially altered. In particular, transduction is required for establishing row 2 identity and for coordinating lengths of stereocilia in a given row.

### Development of stereocilia dimensions in mice

Tilney divided development of the hair bundle into four stages [3]. In stage I (we use Roman numerals here to avoid confusion with row numbers), hair cells undergo their terminal mitoses and begin differentiating. Stereocilia begin to form and elongate in stage II, which also includes initial development of the stereocilia staircase. In mice, stages I and II occur prior to birth; P0.5 mouse IHC bundles in the apex have stereocilia with a discernable staircase (Fig. 1B), which develops before the acquisition of transduction currents. Similar to chick cochlea, stereocilia widening (or narrowing) in mouse IHCs (stage III) occurs between lengthening phases. Changes in diameter of rows 2 and 3 are correlated with the appearance of transduction, while row 1 widening is more protracted and completes several days later in development (Fig. 10A). Finally, there is a late stereocilia elongation phase (stage IV) when mature lengths are attained; during this stage in the mouse, row 1 elongates further but row 2 shortens. In contrast with a previous suggestion [4], we thus find that mouse IHC stereocilia growth segregates into specific lengthening and widening phases, and that these phases align developmentally with the maturation of transduction (Fig. 10A). Tilney’s division of bundle development into four specific stages [3] thus holds true broadly in mouse, albeit with additional complexity.

**Figure 10.**
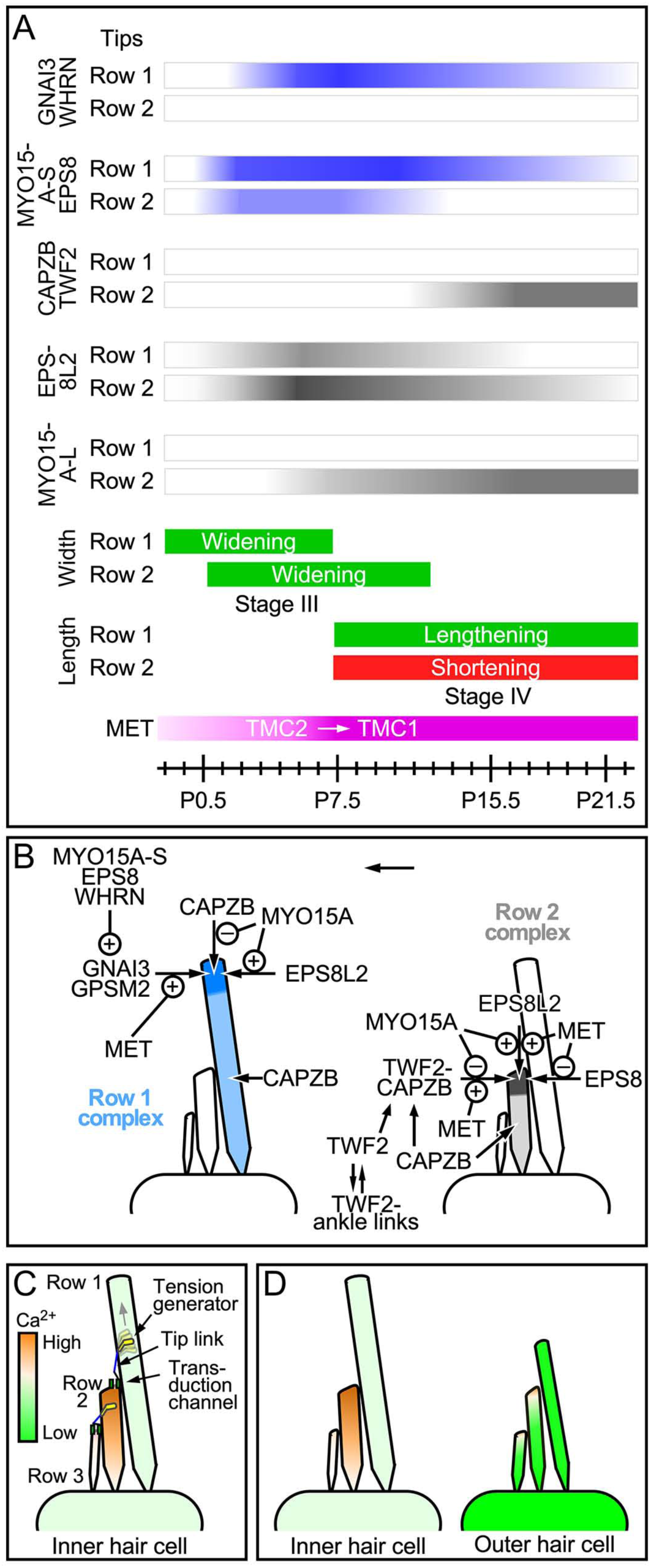
Model for control of rows 1 and 2 dimensions by transduction. ***A***, Expression of row 1 complex proteins (blue bars) and row 2 complex proteins (gray bars) during development. At the bottom, widening occurs as transduction is acquired (stage III), while lengthening (or shortening for row 2) occurs after transduction reaches mature levels (stage IV). ***B***, Row 1 activity (no transduction channels present). Using WHRN as a linker, MYO15A-S and EPS8 deliver GNAI3 and GPSM2 to stereocilia tips, where they control lengthening. Transduction (MET) indirectly affects this complex from accumulating at row 1 tips. MYO15A also stimulates EPS8L2 accumulation at row 1 tips, but because of the presence of EPS8, its levels are reduced compared to those at row 2 tips. Finally, MYO15A prevents CAPZB from accumulating at stereocilia tips. Right, row 2 activity (transduction channels present). Transduction favors the row 2 localization of EPS8L2 over EPS8; the presence of EPS8L2 at stereocilia tips requires MYO15A. CAPZB (and other unknown proteins) controls widening at earlier developmental time points, when TWF2 is sequestered at ankle links. After ankle links disappear (>P10), TWF2 is released, binds to CAPZB, and accumulates at row 2 tips in a transduction-dependent manner. ***C***, Rows 2 and 3 could have different resting Ca^2+^ levels. High motor tension on the row 2>1 tip link should cause high Ca^2+^ levels in row 2. Low motor tension on the row 3>2 tip link should cause low Ca^2+^ levels in row 3. Ca^2+^ in turn could affect stereocilia length and width. ***D***, Differences in Ca^2+^ extrusion and buffering between IHCs and OHCs could produce distinct hair-bundle shapes. Because of high levels of ATP2B2, the plasma-membrane Ca^2+^ ATPase, as well as mobile Ca^2+^ buffers, Ca^2+^ levels should be much lower in OHC stereocilia than in IHC stereocilia.

### Control of stereocilia length and width by transduction

Consistent with the temporal overlap of stereocilia row differentiation with transduction onset during wild-type development, we find that in IHCs of mice lacking transduction throughout early development, rows of stereocilia develop with more uniform widths and heights [19]. Nevertheless, stereocilia still continue to grow to significant lengths, unlike in mutants lacking the row 1 complex proteins MYO15A-S, EPS8, WHRN, GPSM2, or GNAI3; the general mechanisms that stimulate stereocilia growth thus remain present in transduction mutants. Row height is also not completely equalized in transduction mutants, so the basic stereocilia staircase is established using other mechanisms, most likely using links between stereocilia [26, 38]. In transduction mutants, however, the stages of growth that normally follow transduction onset (III and IV) are significantly altered. Stereocilia of transduction mutants failed to undergo the row 2 thickening and row 3 thinning that occurs following transduction onset (stage III), and also did not complete lengthening of row 1 and shortening of row 2 in stage IV. In addition, coordination between stereocilia dimensions of the same row was substantially reduced, especially at older ages, leading to irregular heights and widths. Length and width irregularities were unexpectedly pronounced for row 1 stereocilia in *Tmie* mutants, which was surprising given the absence of transduction channels at row 1 tips but might be explained by redistribution of row 1 and row 2 proteins. Together, these results suggest that transduction plays a key role in determining the stereocilia dimensions of all three rows of stereocilia and in matching heights within a given row.

Pharmacological blockade of transduction channels led to shortened row 2 and 3 stereocilia in cochlear cultures [26]. By contrast, our genetic approach shows that loss of transduction during development is not correlated with shortening of row 2 and 3 stereocilia rows. TMCs are absent from stereocilia not only in *Tmc* DKOs, but also in *Tmie* KOs [23], which raises the possibility that the presence of the TMCs or their interaction partners is required for stereocilia shortening elicited by channel block. In addition, blockade of transduction within a short window of development (≤32 hr), rather than across all developmental stages, may understandably produce a different outcome. As transduction can also regulate the maintenance of uniform heights between stereocilia within a row, the shortening of rows 2 and 3 seen with transduction blockade likely reflects a failure of this maintenance process.

### Control of row identity by transduction

The developmental concentration of EPS8 and MYO15A-S to row 1, correlated with the appearance there of GNAI3 and WHRN, coincides with results of other studies that have shown that GPSM2 and GNAI3 help establish row 1 identity by stabilizing WHRN, MYO15A-S, and EPS8 at tips of this row [10, 11, 14]. The timing of these events suggests that surprisingly, transduction assists in targeting the complex of GNAI3 and GPSM2 complex to row 1 tips, which was supported by our observation that GNAI3 levels at row 1 tips were significantly reduced in transduction mutants, especially as the cells matured (Fig. 5). Decreased GNAI3 at row 1 tips was correlated with redistribution of EPS8, MYO15A-S, and WHRN to all stereocilia tips by P21.5. The complex of GNAI3 and GPSM2 still specifically targeted to row 1 tips during the first week of development in the transduction mutants, albeit at a reduced level; this complex thus initiates differentiation of row 1 early during development but cannot maintain it without transduction. Transduction could plausibly regulate row 1 identity by controlling how MYO15A couples to GNAI3 or WHRN, especially if the complex is formed before it enters the stereocilia.

Distribution of row 2 proteins also depended on transduction. EPS8L2 was present at all stereocilia tips prior to transduction onset, but concentrated at row 2 (and 3) during that row’s shortening period in stage IV. Restriction of EPS8L2 to the shorter rows also coincided with the increased appearance of MYO15A-L at shorter row tips. Both events were disrupted in transduction mutants. As the C-terminal domain in EPS8 that mediates its interaction with the first MYTH-FERM domain of MYO15A [7] is present in EPS8L2 [39], EPS8L2 and EPS8 could each be transported or regulated by MYO15A. CAPZB and TWF2 only appeared at row 2 tips towards the end of the stage IV developmental phase as the cells are nearing maturity, suggesting that these form a transduction-dependent capping complex to maintain row 2 heights after shortening is complete.

Row 2 identity is likely set in part by transduction-dependent alteration of the environment of that row’s tips (Fig. 10B), most likely by elevating Ca^2+^ there—although the physical presence of the transduction proteins might be important as well. In support of the former hypothesis, pharmacological blockade of transduction, which prevents Ca^2+^ entry, initiated accumulation of row 1 proteins at row 2 tips. Ca^2+^ could alter the molecular environment at tips by preventing accumulation of EPS8 and MYO15A-S at row 2 tips or by favoring EPS8L2 accumulation at row 2, which might exclude EPS8 and MYO15A-S. Although EPS8 and EPS8L2 have not been reported to be directly influenced by Ca^2+^, MYO15A might be controlled directly through members of the CALM family that bind to its lever arm [40]. Additional mechanisms could exploit other Ca^2+^-dependent actin regulatory proteins. Notably, mice lacking the Ca^2+^-binding protein CIB2 have a hair-bundle morphology similar to that seen in transduction mutants [41]; CIB2 might mediate Ca^2+^ control of row proteins and, in turn, stereocilia dimensions.

### Two roles for heterodimeric capping protein

CAPZB, one subunit of the heterodimeric capping protein, shows distinct patterns of localization at different developmental times. Before P15.5, CAPZB decorates the outside of stereocilia shafts. As we speculated previously, CAPZB may stabilize actin filaments that are added there and elongate during widening phases [15]. At P15.5 and later, however, CAPZB disappears from stereocilia shafts and appears instead at tips of row 2 stereocilia; its binding partner TWF2 is restricted to the ankle-link region prior to P11.5, but shifts to row 2 tips simultaneously with CAPZB. In inner hair cells, ankle links disappear by P12 [42], leading us to propose that ankle links capture and sequester TWF2 during early stereocilia development, and when the links disappear, TWF2 is released, binding CAPZB and creating a complex that now targets stereocilia tips and controls actin dynamics there (Fig. 10B). Importantly, our experiments demonstrate that transduction is required for localization of CAPZB and TWF2 to row 2 stereocilia tips, although the molecular mechanism is unknown. Late acquisition of CAPZB at row 2 tips may be important for length coordination of this row’s stereocilia.

The absence of MYO15A leads to the loss of EPS8L2 and the accumulation of CAPZB and TWF2 at all stereocilia tips, suggesting that MYO15A controls which actin filament capping molecules are present. In turn, the presence of different cappers may affect length and perhaps width regulation. As *Myo15a*^*sh2*^ mutants share a similar morphological phenotype with other row 1 protein mutants, early accumulation of CAPZB and TWF2 at all stereocilia tips likely occurs in those mutants as well. Indeed, other studies have observed TWF2 at all stereocilia tips in *Myo15a, Whrn, Gpsm2*, and *Gnai3* mutant IHC [10, 43, 44]. The anomalous early presence of the CAPZB-TWF2 capping complex at all tips in these mutants may explain in part why their stereocilia remain uniformly short. Together these results suggest that transduction helps to time the capping of actin filaments.

### Differential control of row dimensions by transduction

A notable paradox is why transduction controls row 2 and row 3, each with transduction channels, in distinct ways. While each row shrinks after transduction appears during development, row 2 widens over the first few days while row 3 narrows. Differences in transduction-channel open probability and hence Ca^2+^ entry can explain why the two rows grow differentially (Fig. 10C). Because it is present in channel-lacking row 1, the hypothesized tensioning motor at the upper end of the tip link—which controls the row 2 transduction channel [1]—should see a lower Ca^2+^ concentration than the motor controlling the row 3 channel [45]. In turn, low Ca^2+^ in row 1 will increase motor force production and increase tip-link tension, leading to a high open probability for the row 2 channel [46]. By contrast, Ca^2+^ can enter at row 2 tips and diffuse down to the anchor for the row 3 channel’s tip link; the high Ca^2+^ affecting its tip-link motor (within row 2) will reduce force production and decrease open probability of the row 3 channel. Thus resting Ca^2+^ should be high in row 2 but low in row 3 (Fig. 10C). Accurate differential measurement of Ca^2+^ in each row could provide support for this hypothesis but will be technically difficult to carry out.

A related question poses why the stereocilia dimensions of inner and outer hair cells are so different. Differential control of Ca^2+^ levels within stereocilia could be responsible for these morphological differences. Compared to OHCs, IHCs have much lower concentrations of mobile Ca^2+^ buffers [47] and a much lower density of the principal Ca^2+^ pump ATP2B2 [48, 49]. Ca^2+^ entering through IHC transduction channels will thus reach higher levels and extend further longitudinally in IHCs than in OHCs. Given that the control of actin-core dimensions by transduction is likely through entering Ca^2+^, the highly asymmetric stereocilia dimensions of IHC hair bundles likely result from a greater impact of entering Ca^2+^ (Fig. 10D).

## Conclusions

Transduction is essential to establish the normal dimensions of row 2 stereocilia and for the acquisition of that row’s molecular identity. Transduction also plays an unexpected role in the regulation and maintenance of row 1 stereocilia dimensions, despite the lack of transduction at row 1 tips. The loss of transduction normalizes the molecular identity of stereocilia rows so that they are essentially the same, most similar to the identity of wild-type row 1. Transduction also regulates capping complexes, which contribute to maintenance of stereocilia dimensions into adulthood. The maturation of transduction is thus tightly interconnected to regulation of stereocilia dimensions, allowing the functional development of the hair bundle to guide its morphological development.

## Acknowledgements

We greatly appreciate excellent mouse husbandry from Jennifer Goldsmith and Ruby Larisch. We thank Andy Griffith for the *Tmc1*^*KO*^ and *Tmc2*^*KO*^ mouse lines, Uli Müller for the *Tmie*^*KO*^ mouse line, Gregory Frolenkov for the *Myo15a*^*sh2*^ mouse line, and Stefan Heller for the *Twf2*^*KO*^ mouse line. Tom Friedman generously provided anti-MYO15A antibodies TF1, PB888, and PB48; Uli Müller provided WHRN antibodies; Stefan Heller provided TWF2 antibodies. We carried out confocal microscopy at the OHSU Advanced Light Microscopy Core @ The Jungers Center (P30 NS061800 provided support for imaging); electron microscopy was performed at the OHSU Multiscale Microscopy Core. JFK was supported by NIH grant R03 DC014544. JEB was supported by startup funds from the University of Florida. PGBG was supported by NIH grants R01 DC002368 and R01 DC011034.

## Author contributions

JK designed experiments, carried out most of the confocal microscopy, analyzed and interpreted data, and participated in manuscript writing; PC conducted stereocilia length and width measurements and electron microscopy and edited the manuscript; RD assisted with electron microscopy and edited the manuscript; DC carried out statistical analyses and edited the manuscript; JB designed experiments, assisted with experiment design and analysis, and edited the manuscript; and PBG assisted with experiment design and analysis, prepared the figures, and wrote the manuscript.

## Experimental Procedures

### Mice

All animal procedures were approved by the Institutional Animal Care and Use Committee (IACUC) at Oregon Health & Science University (protocol IP00000714). Mouse pups were assumed to be born at midnight, so animals used on the first day are referred to as P0.5. Both male and female pups were used. For developmental analysis and cochlear culturing, we used C57BL/6J mice (RRID:IMSR_JAX:000664, Jackson Laboratories, Bar Harbor, ME). The *Tmc1*^*KO*^ and *Tmc2*^*KO*^ lines have been described [20]; animals have been maintained on a C57BL/6J background. Most experiments used animals from *Tmc1*^*KO/KO*^;*Tmc2*^*KO*^*/+* (female) x *Tmc1*^*KO/KO*^;*Tmc2*^*KO/KO*^ (male) crosses. Some experiments used animals from *Tmc1*^*KO/KO*^;*Tmc2*^*KO*^*/+* (female) x *Tmc1*^*KO*^*/+;Tmc2*^*KO/KO*^ (male) crosses to examine both single and double knockout littermates. Mice were genotyped using previously described primers [20]. The *Tmie*^*KO*^ line has also been described [24] and was maintained on C57BL/6J background. Experiments used animals from *Tmie*^*KO*^*/+* (female) x *Tmie*^*KO/KO*^ (male) crosses. Mice were genotyped using previously described primers (Zhao et al., 2014). The *Myo15a*^*sh2*^ mouse line has been previously described [12] and was obtained from Gregory Frolenkov; we maintained these mice on their mixed background. Genotyping was carried out by Transnetyx (Cordova, TN) using the protocol for Jackson Laboratories stock Cat#000109. Apical, mid, and basal sections of the cochlea refer to the apical-most ∼1/3, the middle ∼1/3, and the basal ∼1/3 of the cochlea.

### Statistical analysis

Pairwise comparisons (e.g., row 1 vs row 2 at a specific developmental time) used the Student’s t-test (unpaired samples, two-tailed). Four-way comparisons between row 1 and row 2 from control and mutant cochleas used linear mixed-effects models with y-transformed in the log scale. For each comparison, cochleas from 2-3 mice were used, and 1-3 images were used from each cochlea. A random intercept was included in the model to account for potential correlations among intensity values within cochleas and images. Values of 0 were replaced with 0.0001 for the log transformation of y. All computations were done by the lmerTest R package [50] in the R statistical language (http://www.r-project.org). All computations were done by the lmerTest R package [50] in the R statistical language (http://www.r-project.org).

### Immunofluorescence microscopy

Inner ears were isolated from C57BL/6J mice or mutant mice littermates at the indicated ages and dissected in cold Hank’s balanced salt solution (Cat#14025076, Thermo Fisher Scientific, Waltham, MA) supplemented with 5 mM HEPES, pH 7.4 (dissection buffer). Images of tip proteins labeled “P21.5” were dissected from weaning-age mice between P19.5 and P23.5; images for measurement of stereocilia dimensions were from the exact ages marked on figures. For all organs, small openings were made within the periotic bones to allow perfusion of the fixative. For measurement analysis of C57BL/6J or mutant mice using only phalloidin (Figs. 1-2), ears were fixed in 4% formaldehyde (Cat#1570, Electron Microscopy Sciences, Hatfield, PA) in 1x PBS for 12-24 hours at 4°C. Ears were washed in PBS, then cochleas were dissected out from the periotic bone and the lateral wall was removed. Cochleas were permeabilized in 0.5% Triton X-100 in 1x PBS for 10 minutes at room temperature, then incubated with phalloidin (0.4 U/ml Alexa Fluor 488 phalloidin; Cat#A12379, Thermo Fisher Scientific) in 1x PBS for 3-4 hours at room temperature. Organs were washed three time in PBS for 5 min per wash and then mounted in Vectashield (Cat#H-1000, Vector Laboratories, Burlingame, CA).

For immunofluorescence using antibodies against row 1 or 2 proteins (Figs. 3-4, 7-9), ears were fixed in 4% formaldehyde (Cat#1570, Electron Microscopy Sciences) in dissection buffer for 20-60 min at room temperature. Length of fixation depended on the primary antibody used in an experiment; we found that row 2 protein antibodies (EPS8L2, CAPZB, TWF2) required that fixation last no more than 20 min, whereas the other antibodies were not sensitive to fixative duration. Ears were washed twice in PBS, then cochleas were dissected from periotic bones and the lateral wall was removed. Cochleas were permeabilized in 0.2% Triton X-100 in 1x PBS for 10 min and blocked in 5% normal donkey serum (Cat#017-000-121, Jackson ImmunoResearch, West Grove, PA) diluted in 1x PBS (blocking buffer) for 1 hr at room temperature. Organs were incubated overnight at 4°C with primary antibodies in blocking buffer at the dilutions indicated in Table S1 and then washed three times in 1x PBS. Tissue was then incubated with secondary antibodies, which were either 2 µg/ml donkey anti-mouse Alexa Fluor 488 (Cat#A21202, Thermo Fisher Scientific), 2 µg/ml donkey anti-rabbit Alexa Fluor 488 (Cat#A21206, Thermo Fisher Scientific); 2 µg/ml donkey anti-mouse Alexa Fluor 568 (Cat#A10037, Thermo Fisher Scientific), or 2 µg/ml donkey anti-rabbit Alexa Fluor 568 (Cat#A10042; Thermo Fisher Scientific); 1 U/ml CF405 phalloidin (Cat#00034, Biotium, Fremont, CA) was also included for the 3-4 hr room temperature treatment. When only one primary antibody was used, organs were incubated as above with Alexa Fluor 488 secondary antibody and 0.4 U/ml CF568 phalloidin (Cat#00044, Biotium). Tissue was washed three times in PBS and mounted on a glass slide in ∼50 µl of Vectashield and covered with a #1.5 thickness 22 x 22 mm cover glass (Cat#2850-22, Corning, Corning, NY).

Organs were imaged using a 63x, 1.4 NA Plan-Apochromat objective on a Zeiss Elyra PS.1/LSM710 system equipped with an Airyscan detector and ZEN 2012 (black edition, 64-bit software; Zeiss, Oberkochen, Germany) acquisition software. Settings for x-y pixel resolution, z-spacing, as well as pinhole diameter and grid selection, were set according to software-suggested settings for optimal Nyquist-based resolution. Raw data processing for Airyscan-acquired images was performed using manufacturer-implemented automated settings. For each antibody, 2-4 images were acquired from 1-2 cochlea per genotype per age for each experiment, and experiments were repeated at least twice. Ears from control and mutant littermates and from different ages of C57BL/6J mice were stained and imaged on the same days for each experiment to limit variability. During image acquisition, the gain and laser settings for the antibody and phalloidin signals were adjusted to reveal the staining pattern in control samples, and the corresponding DKO or KO samples used the same settings. Image acquisition parameters and display adjustments were kept constant across ages and genotypes for every antibody/fluorophore combination. Display adjustments in brightness and contrast and reslices and/or average Z-projections were made in Fiji/ImageJ software.

For point spread function (PSF) measurements, 100 nm diameter TetraSpeck microspheres (Cat#T7279, Thermo Fisher Scientific) were dried onto a coverslip (Cat#2850-22, Corning), and then mounted onto a slide with Vectashield. A z-stack was acquired around the center of the beads using the parameters used for our experiments. Single beads were autodetected within the stack and channel-specific point spread functions were calculated using the experimental PSF dialog in ZEN software using default parameters. The PSF profile was exported as a text file, then fit with a single Gaussian. The full width at half-maximum (FWHM) of the PSF was calculated from the fit.

### Scanning electron microscopy

For examining hair-bundle development, C57BL/6J mice at P0.5, P3.5, and P7.5 were used. After post-fixation, tissues were processed for SEM by the OTOTO (osmium-thiocarbohydrazide-osmium) method. Tissues were incubated for 1 hour in 1% OsO_4_, washed in distilled water, and further incubated in 1% thiocarbohydrazide for 20 min. This incubation sequence was repeated for a total of three times. Next, the cochleas were dehydrated with graded ethanol and critical-point dried using liquid CO_2_. Finally, the samples were mounted on aluminum stubs using colloidal silver and imaged using a Helios Nanolab 660 DualBeam Microscope from FEI (Hillsboro, OR).

For comparing controls with mutants, periotic bones with cochleas were dissected in Leibowitz’s L-15 medium (Invitrogen, Carlsbad, CA) from P8.5 littermates from *Tmie, Tmc1;Tmc2*, and *Myo15a*^*sh2*^ crosses. An age-matched C57BL/6J control group was also included. After isolating the periotic bone, several small holes were made to provide access for fixative solutions; encapsulated cochleas were fixed for an hour in 2.5% glutaraldehyde in 0.1 M cacodylate buffer supplemented with 2 mM CaCl_2_. Next, cochleas were washed with distilled water and the cochlear sensory epithelium was dissected out; the tectorial membrane was manually removed. The cochlear tissues were then transferred to scintillation vials and dehydrated in a series of ethanol and critical-point dried using liquid CO_2_. Samples were immobilized on aluminum specimen holders using a carbon tape and sputter coated with 3-4 nm of platinum. Samples were imaged using the Helios scanning electron microscope.

### Quantitation of stereocilia length and width

For analyzing length and width of apical IHC stereocilia during development, z-stack images of phalloidin-stained IHC stereocilia from cochleas of C57BL/6J mice were obtained using identical image acquisition parameters. Although image-scanning microscopy with Airyscan detection measures length and width convolved with the objective’s point-spread function, and not absolute dimensions, our measurements nonetheless provide a valuable view of developmental changes from the measured relative changes in dimensions. Moreover, the large size of IHC stereocilia substantially improved accuracy of our measurements, especially width, reducing the relative uncertainty introduced by the point-spread function. By contrast, accurate stereocilia measurements with scanning electron microscopy require image acquisition with at least two tilt values [26]. Additionally, these samples undergo shrinkage during the dehydration and drying steps, which alters stereocilia dimensions, and stereocilia surfaces rather than boundaries of the actin core are measured. With z-stacks acquired through hair bundles, we can reconstruct the actin cores of all stereocilia, including those that might be obscured by imaging from the bundle’s side. While an ideal technique for measuring actin core dimensions may be focused ion beam scanning electron microscopy [51], throughput with that technique is very low.

Images were acquired on a ZEISS Elyra PS.1/LSM 710 Airyscan combination setup, using a 63x 1.4 NA Plan-Apochromat or a 100x 1.46 NA alpha Plan-Apochromat lens, integrated under ZEN 2.1 software. Raw data processing for Airyscan-acquired images was performed using manufacturer-implemented automated settings. At least 2-4 cochleas were used for developmental ages E18.5 (embryonic), P0.5, P4.5, P7.5, P10.5, P12.5, P15.5, P19.5 (post-natal). For each developmental age, images containing hair bundles that were upright were selected for generating x-z reslices (Fig. S1C) of stereocilia with the ‘Reslice’ function in Fiji. Reslices were chosen with views of stereocilia rows 1 and 2; we chose central stereocilia in most bundles. Stereocilia lengths were measured individually in Fiji by manually drawing a vertical line from the top of a stereocilium to the point of insertion in the cuticular plate. For measuring stereocilia widths, horizontal lines were manually drawn at 50% of the bundle height for both rows 1 and 2. Dimensions of mid-cochlea and basal IHC stereocilia were also measured using the same strategy. Stereocilia dimensions were similarly measured in *Tmie, Tmc1;Tmc2* and *Myo15a*^*sh2*^ mutant mice at P7.5 and P21.5. Original quantitation data of stereocilia length and width during C57BL/6 development from Fig. 1 are available from figshare.com using the identifier 10.6084/m9.figshare.9275156. Original quantitation data of stereocilia length and width of *Tmc* DKO, *Tmie* KO, and *sh2* KO mice from Figs. 2 and S3 are available from figshare.com using the identifier 10.6084/m9.figshare.9275168.

### Quantitation of proteins at rows 1 and 2 tips

We estimated the total immunofluorescence signal in the top 1-2 µm of each tip. Airyscan z-stacks were imported into Fiji, which was used for all analysis steps. For analysis of P0.5 and P7.5 C57BL/6J mice and P7.5 mutant mice, average Z-projections of Airyscan stacks were made that included row 1 and row 2 tips in the same projection. For analysis of P15.5 and P21.5 C57BL/6J mice, separate Z-projections were made for row 1 and row 2 tips, using the same number of projected x-y slices per row. For analysis of P21.5 mutant mice, due to increased height variability, Regions of Interest (ROIs) were selected at row 1 or row 2 tips within individual x-y slices from the z-stacks. Images were kept as multi-channel stacks and phalloidin was used to guide selection at the tips of each row. Regions of interest (ROIs) used circles that encompassed most of each tip. Area and average intensity measurements were made from ten or more row 1 tips and ten or more row 2 tips, and blank measurements were taken from volumes outside the stereocilia and above the epithelium. The total signal (area times average intensity) in each stereocilia tip volume was calculated after subtracting the background. For each hair bundle, the average signal in rows 1 and 2 was calculated, which was used to normalize the individual row 1 and row 2 stereocilia tip signals within each bundle. The average signal row 1 and 2 signal for each bundle was also used for comparison of expression levels at different developmental times or between control and mutant mice. For developmental comparisons, the average signal in rows 1 and 2 for each bundle was normalized to the highest average value for the protein’s developmental series. For comparisons between transduction-mutant genotypes, the average signal for control bundles on a given experimental day was used to normalize all genotypes, allowing comparison of controls and mutants with identical acquisition parameters. Data shown represents analysis of at least 4 images per condition, from at least two separate experiments. Original quantitation data of C57BL/6 development from Figs. 3 and 7 are available from figshare.com using the identifier 10.6084/m9.figshare.9275141. Original quantitation data of transduction mutants from Figs. 5, 8, and S4 are available from figshare.com using the identifier 10.6084/m9.figshare.9253217.

### Cochlea culture and tubocurarine block

The channel-block methods were adapted from previous experiments [26]. Organ of Corti explants were isolated from male and female wild-type C57BL/6J mice at P4. The explants were trimmed using a sapphire knife (World Precision Instruments, Sarasota, FL) to include only the middle turn and were lightly adhered to the bottom of a 35 x 10 mm Easy Grip petri dish (Cat#351008, Falcon-Corning). Explants were cultured at 37°C and 5% CO_2_ in DMEM (Invitrogen) supplemented with 7% fetal bovine serum (FBS, Gibco) and 10 mg/ml ampicillin. For channel-blocking experiments, explants were cultured for 24 hr in FBS/ampicillin-supplemented DMEM at 37°C and 5% CO_2_ in the presence or absence of 30 µM tubocurarine (Cat#T2379, Sigma-Aldrich, St. Louis, MO). Following the incubation period, explants were transferred to a 24-well plate containing a 4% formaldehyde solution (Cat#1570, Electron Microscopy Sciences) and fixed for 20 minutes at room temperature and then washed twice in 1x PBS. Explants were stained using EPS8 and MYO15A-S antibodies using the immunofluorescence and Airyscan imaging protocols outlined above.

For FM1-43 labeling, following the 24 hr culture period, explants were incubated for 30 sec in ice-cold Ca^2+^-containing standard HBSS (Cat#14025076, Thermo Fisher Scientific) supplemented with 6 µM FM1-43FX (Cat#F35355, Thermo Fisher Scientific) in the absence or presence of 30 µM tubocurarine. After rinsing thoroughly with HBSS, explants were fixed in 4% formaldehyde solution for 20 min. The samples were rinsed with HBSS, mounted on a slide with Vectashield (Vector Labs) and imaged soon after fixation. Confocal images of FM1-43 treated explants were imaged at room temperature with an integrated photomultiplier tube on an LSM 710 (Zeiss) confocal microscope with a 63x PlanApochromat 1.4 NA oil objective. Images were acquired using Zen 2.1 (Zeiss) Software and processed using Fiji Software. Image acquisition settings and processing steps were kept identical between all experimental conditions. Analysis represents data from at least three separate culture experiments performed on different days.

## Abbreviations

DKO: double knockout;
FWHM: full width at half-maximum;
IHC: inner hair cell;
KO: knockout;
MET: mechanoelectrical transduction;
OHC: outer hair cell.

## Supplemental Figures and Captions

**Figure S1 related to Figure 1.**
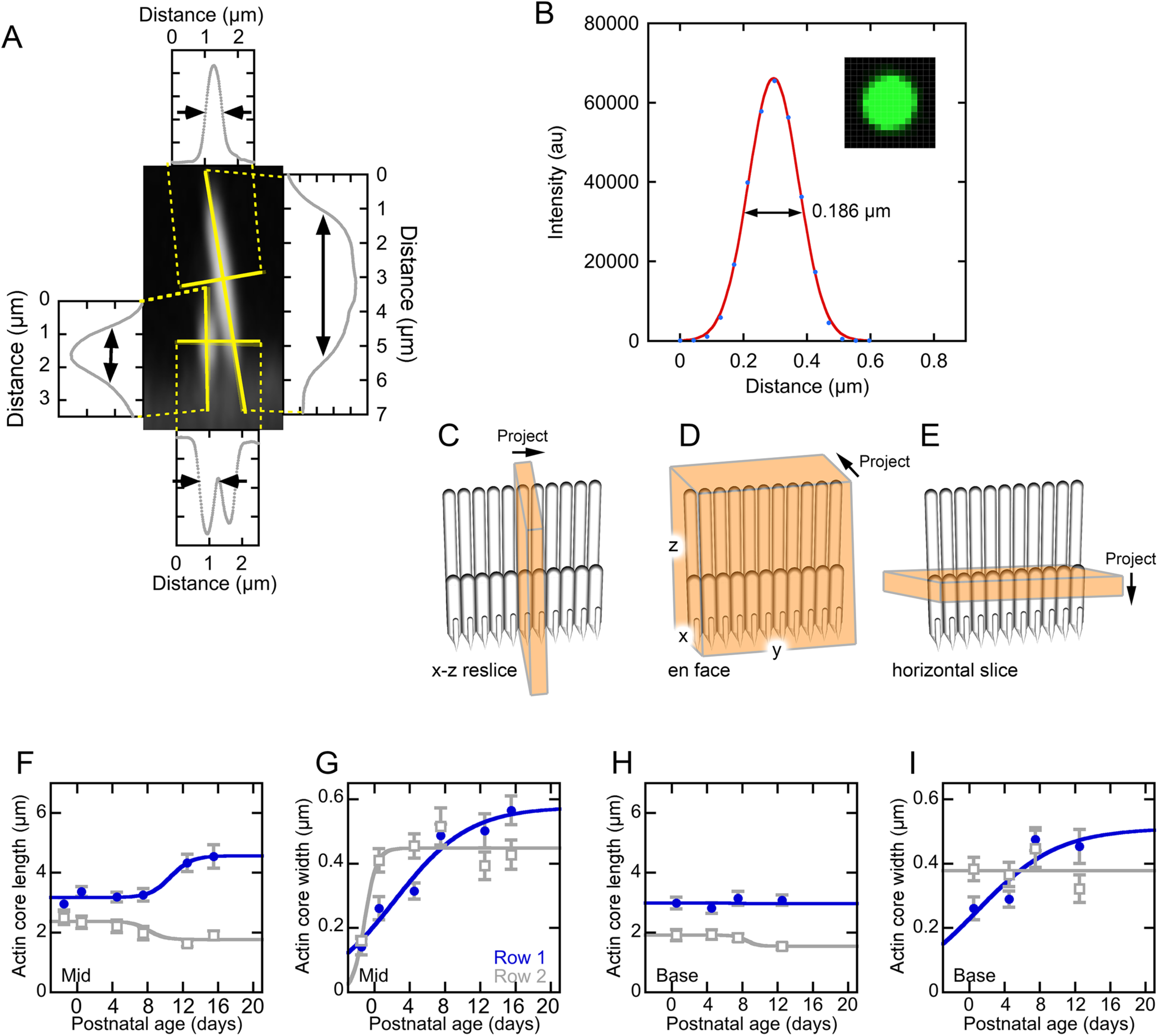
Inner hair cell measurements. ***A***, Length and width measurements. The Plot Profile tool of Fiji was used to calculate longitudinal and cross-sectional profiles on a reslice image (solid yellow lines). Examples of resulting profiles are shown, connected by dashed lines. The unlabeled axis in each plot is fluorescence (arbitrary units). ***B***, Determination of spatial resolution from the PSF. The PSF spatial extent as calculated from the Zen analysis software is plotted and fit with a single Gaussian. The FWHM of 186 nm is indicated. Inset, image of 100 nm TetraSpeck microsphere. ***C-E***, Orientation and generation of images used here. ***C***, x-z reslice. Fiji’s Reslice tool was used to create a single x-z slice or a stack of x-z slices; the stack was projected as shown. Hair bundles are seen in profile. ***D***, en face view. A stack of x-z slices was created, which was projected in y. Bundles are seen with all rows superimposed on each other. ***E***, Horizontal slice. A stack of x-y slices was created, which was projected in z. Cross-sections of all stereocilia in the x-y plane are seen. ***F-I***, Measurements of length and width for mid-cochlea (F, G) and basal cochlea (H, I) inner hair cells during development. Apparent actin core lengths are plotted (F, H) for rows 1 (blue in all plots) and 2 (gray in all plots) during IHC development. Apparent actin core widths are plotted (G, I) for rows 1 and 2 of IHCs during development. Logistic equation fits to each data set.

**Figure S2 related to Figure 2.**
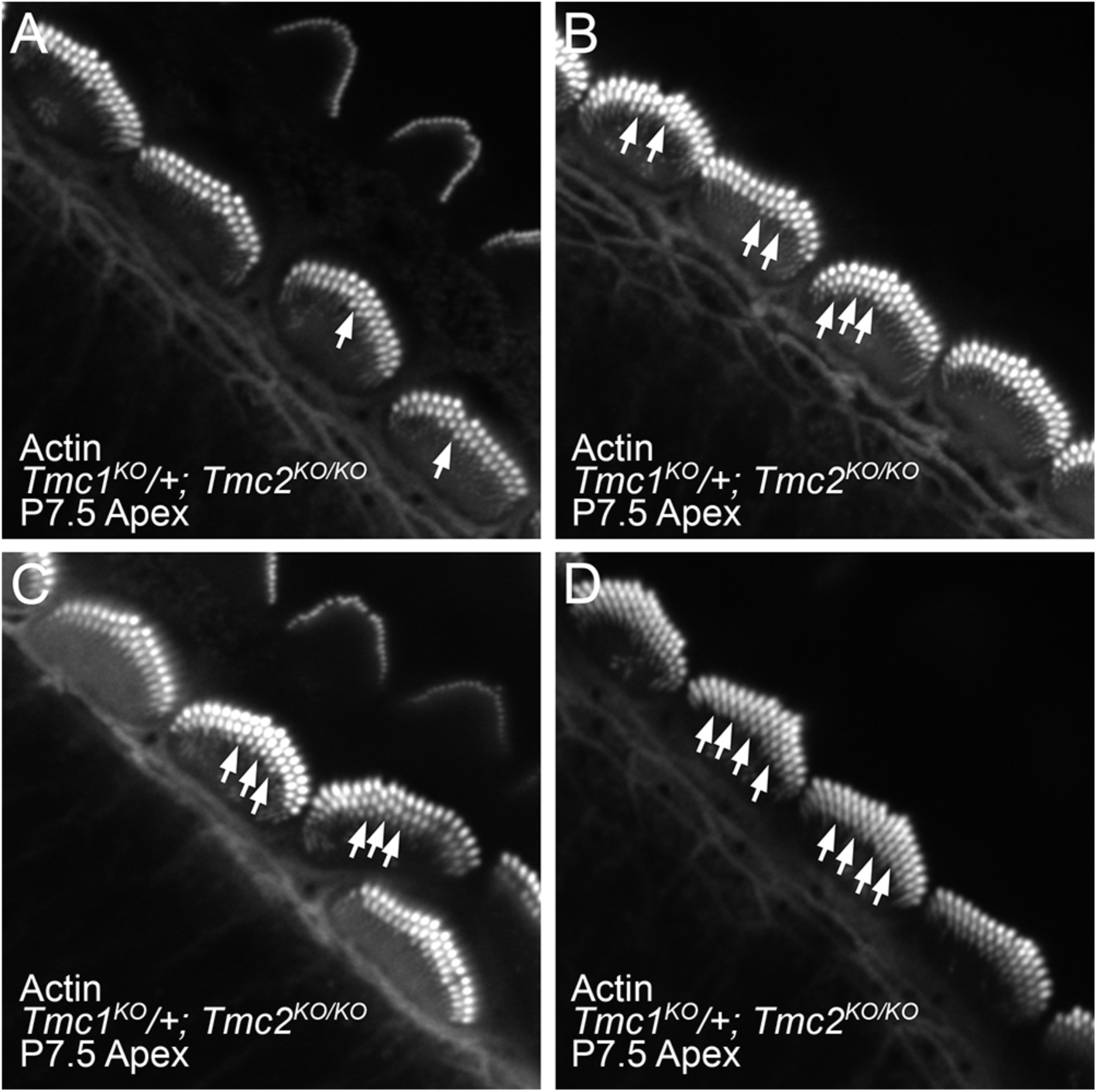
Intermediate phenotype of *Tmc1*^*KO*^/+; *Tmc2*^*KO/KO*^ hair bundles. ***A-D***, Four examples of *Tmc1*^*KO*^/+; *Tmc2*^*KO/KO*^ (P7.5 apex) IHCs labeled for actin (gray). All samples have mostly upright bundles. Arrows indicate thick row 3 stereocilia. All row 3 stereocilia are thick in D. All A-D panels are 29 µm wide.

**Figure S3 related to Figure 2.**
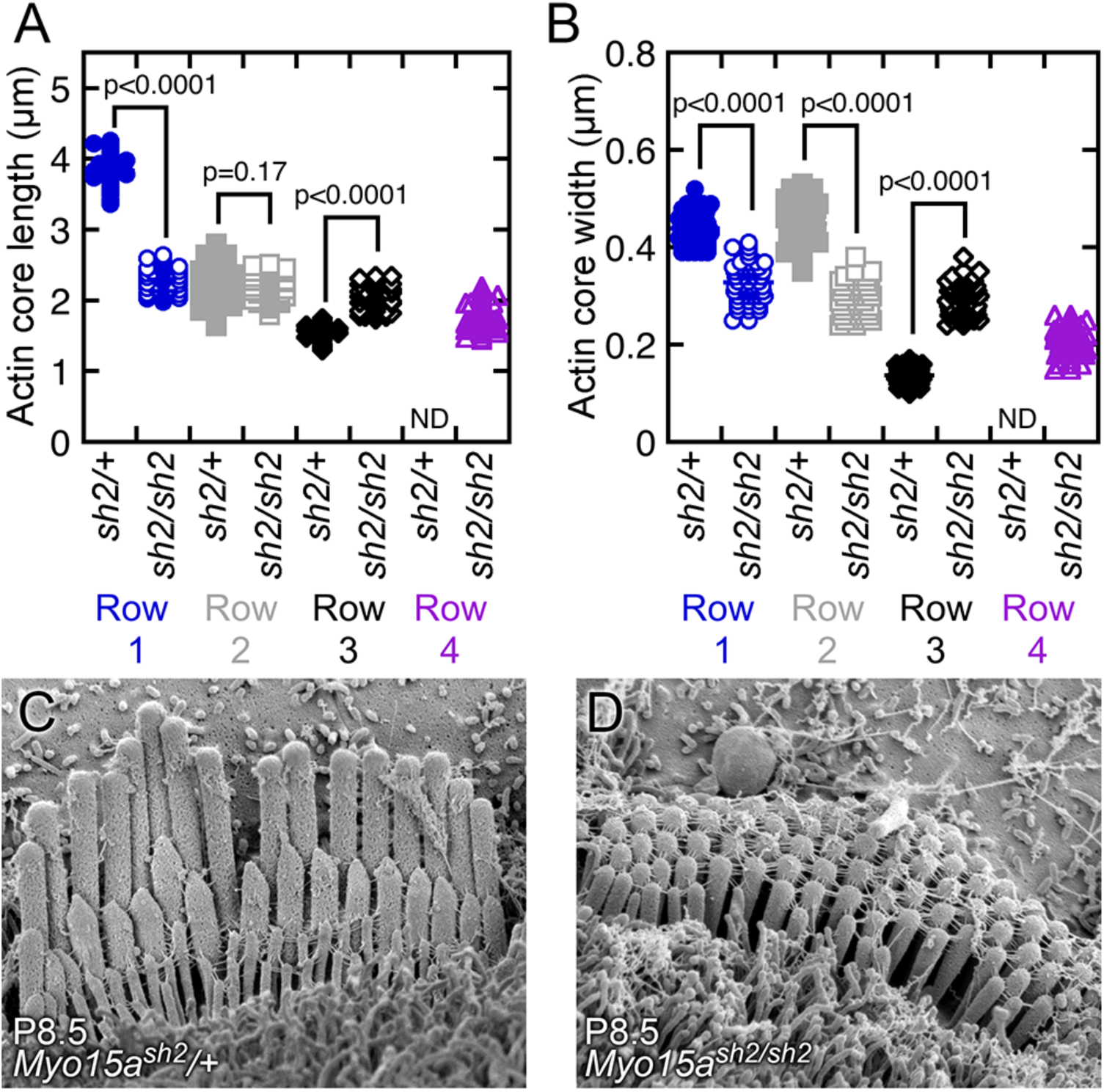
Dimensions of *Myo15a*^*sh2*^ stereocilia. ***A-B***, Quantitation of length (A) and width (B) of P8.5 IHC stereocilia from *Myo15a*^*sh2*^*/+* control (*sh2/+*) and *Myo15a*^*sh2/sh2*^ homozygous mutant (*sh2/sh2*) mice. Pairwise comparisons used t-tests (row 1, n=56-65; row 2, n=56-65; row 3, n=24-65; row 4, n=53). ***C-D***, Scanning electron microscopy images of *Myo15a*^*sh2*^*/+* (C) and *Myo15a*^*sh2/sh2*^ (D) genotypes from apical cochlea at P8.5 (abbreviated respectively *sh2/+* and *sh2/sh2*). Panels are 5.2 µm wide.

**Figure S4 related to Figure 3.**
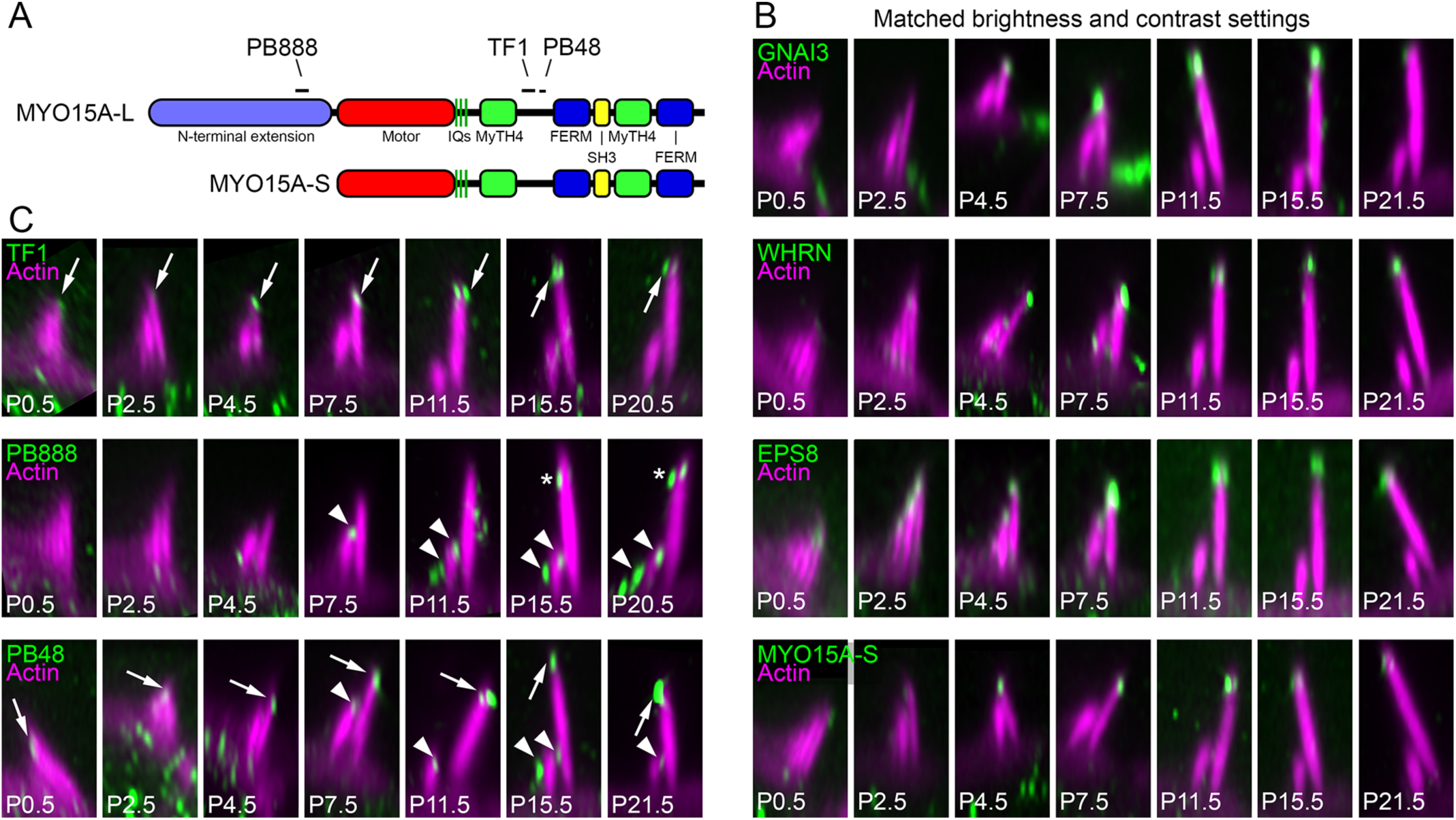
Developmental changes in inner hair cells. ***A***, Domain structure of MYO15A-L and MYO15A-S proteins. Note that while PB888 recognizes an epitope in the unique N-terminal extension of MYO15A-L, both TF1 and PB48 were raised against epitopes that should be shared by MYO15A-L and MYO15A-S. ***B***, Specificity of the TF1 antibody for MYO15A-S. Labeling for TF1, PB888, and PB48 antibodies in apical IHCs over development. Arrows indicate row 1 tip staining, while arrowheads indicate rows 2 and 3 staining. Note that the PB888, specific for a sequence in MYO15A-L, labels row 2 and 3 with occasional row 1 labeling in older cochleas. PB48, which recognizes an epitope shared by MYO15A-L and MYO15A-S, labels all three rows of stereocilia. By contrast, TF1, while directed against an epitope shared by MYO15A-L and MYO15A-S, only recognizes MYO15A at row 1 stereocilia tips. All panels are 4 µm wide. ***C***, Localization of proteins specific for row 1 stereocilia tips in IHC hair bundles during postnatal development. All images are x-z reslices from z-stacks. Brightness and contrast settings in Fiji were identical for each set of proteins, allowing comparisons of signal intensities during development. All panels are 4 x 7.5 µm; in some cases, images were expanded with black background to fill the panel.

**Figure S5 related to Figure 4.**
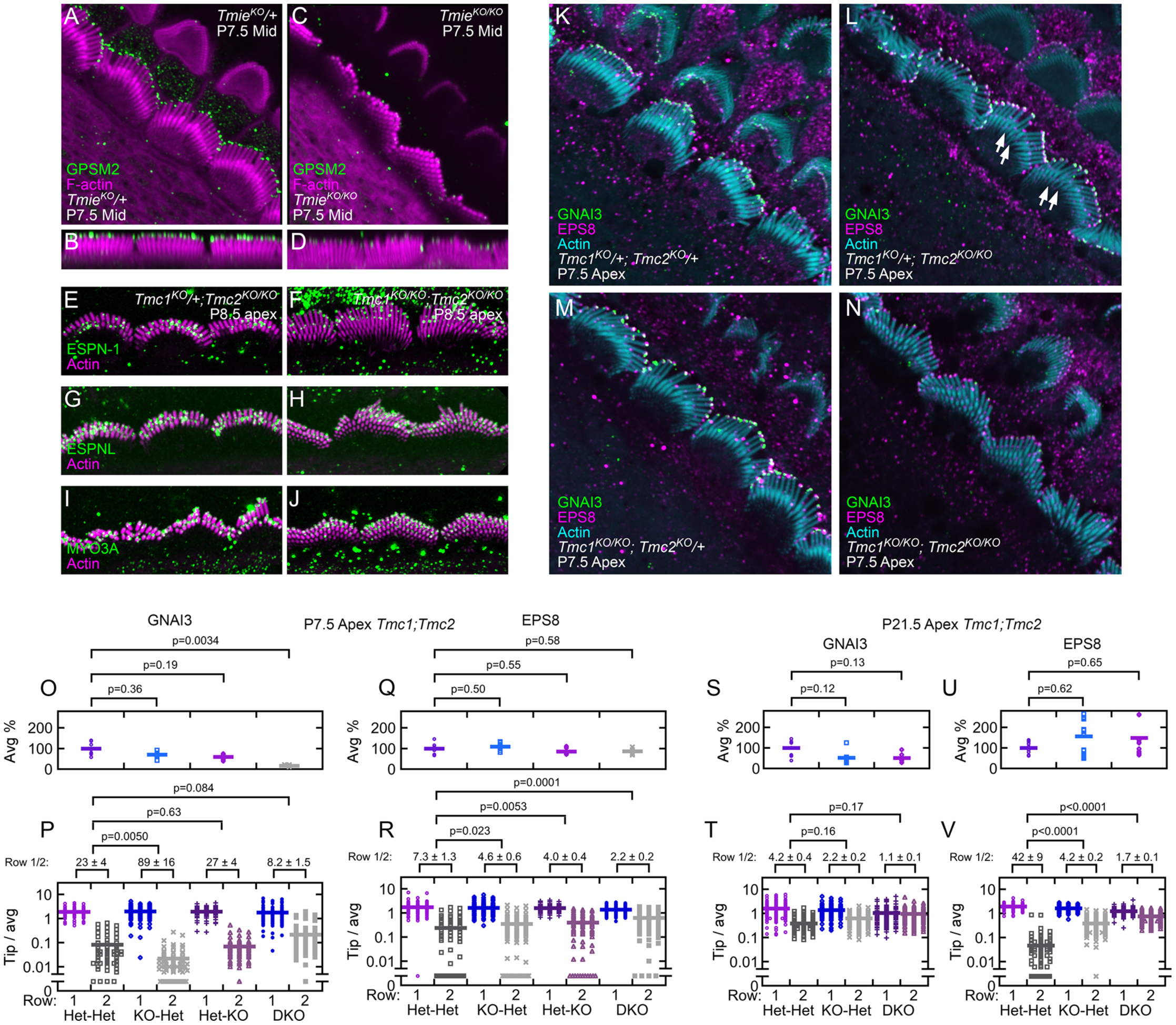
Morphology and row 1 proteins in mutant mice. ***A-D***, Localization of GPSM2 in *Tmie* control (A-B) and *Tmie* KO (C-D) samples. Each pair of control and KO panels were from littermates, with identical sample preparation and confocal imaging parameters. The gain of the antibody signal was adjusted linearly to reveal the staining pattern in each control sample, and the corresponding KO sample used the same gain settings. A and C are x-y sections through a z-stack; B and D are en face views of bundles. All A-D panels are 25 µm wide. ***E-J***, ESPN-1 (E-F), ESPNL (G-H), and MYO3A (I-J) localization in P8.5 apical *Tmc* DKO IHCs. Left panels, *Tmc1*^*KO/KO*^; *Tmc2*^*KO*^/+ controls. Right panels, *Tmc1*^*KO/KO*^; *Tmc2*^*KO/KO*^ DKOs. Each pair of control and DKO panels were from littermates, with identical sample preparation and confocal imaging parameters. The gain of the antibody signal was adjusted linearly to reveal the staining pattern in each control sample, and the corresponding DKO sample used the same gain settings. ***K-N***, *Tmc* genotypes. P7.5 apex IHCs labeled for GNAI3 (green), EPS8 (magenta), and actin (cyan). Arrows indicate thick row 3 stereocilia. All K-N panels are 29 x 29 µm. ***O-R***, Quantitation of row 1 proteins in *Tmc1;Tmc2* mutants at P7.5. Measurements of average signal for all rows (O, Q) and individual tip signal over row 1 plus row 2 average signal (P, R) for GNAI3 (O-P) and EPS8 (Q-R). ***S-V***, Quantitation of row 1 proteins in *Tmc1;Tmc2* mutants at P21.5. Measurements of average signal (S, U) and row over row average (T, V) for GNAI3 (S-T) and EPS8 (U-V). *Tmc1;Tmc2* genotypes: Het-Het, *Tmc1*^*KO*^/+;*Tmc2*^*KO*^/+; KO-Het, *Tmc1*^*KO/KO*^;*Tmc2*^*KO*^/+; Het-KO, *Tmc1*^*KO*^/+;*Tmc2*^*KO/KO*^; DKO, *Tmc1*^*KO/KO*^;*Tmc2*^*KO/KO*^.

**Figure S6 related to Figure 7.**
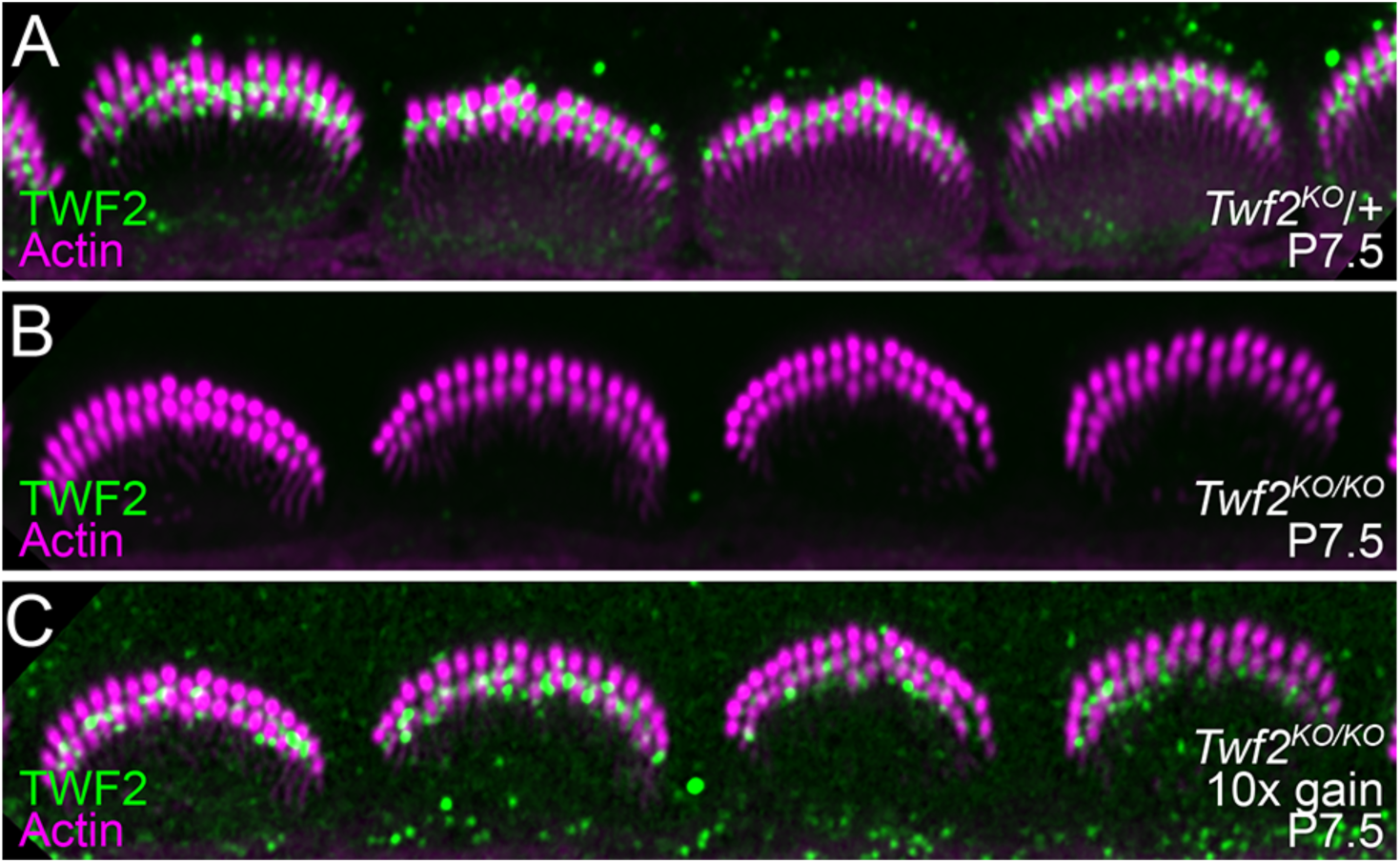
TWF2 staining in *Twf2*^*KO/KO*^ cochleas. ***A***, TWF2 (green) and F-actin (magenta) labeling of *Twf2*^*KO*^*/+* heterozygotes; images show apical IHCs at P7.5. ***B***, Labeling of *Twf2*^*KO/KO*^ homozygotes. ***C***, Labeling of *Twf2*^*KO/KO*^ homozygotes with gain increased 10-fold. Note that the TWF2 antibody crossreacts somewhat with TWF1 [1], which is also localized in stereocilia [2]. Panels are 40 µm wide.

**Figure S7 related to Figure 8.**
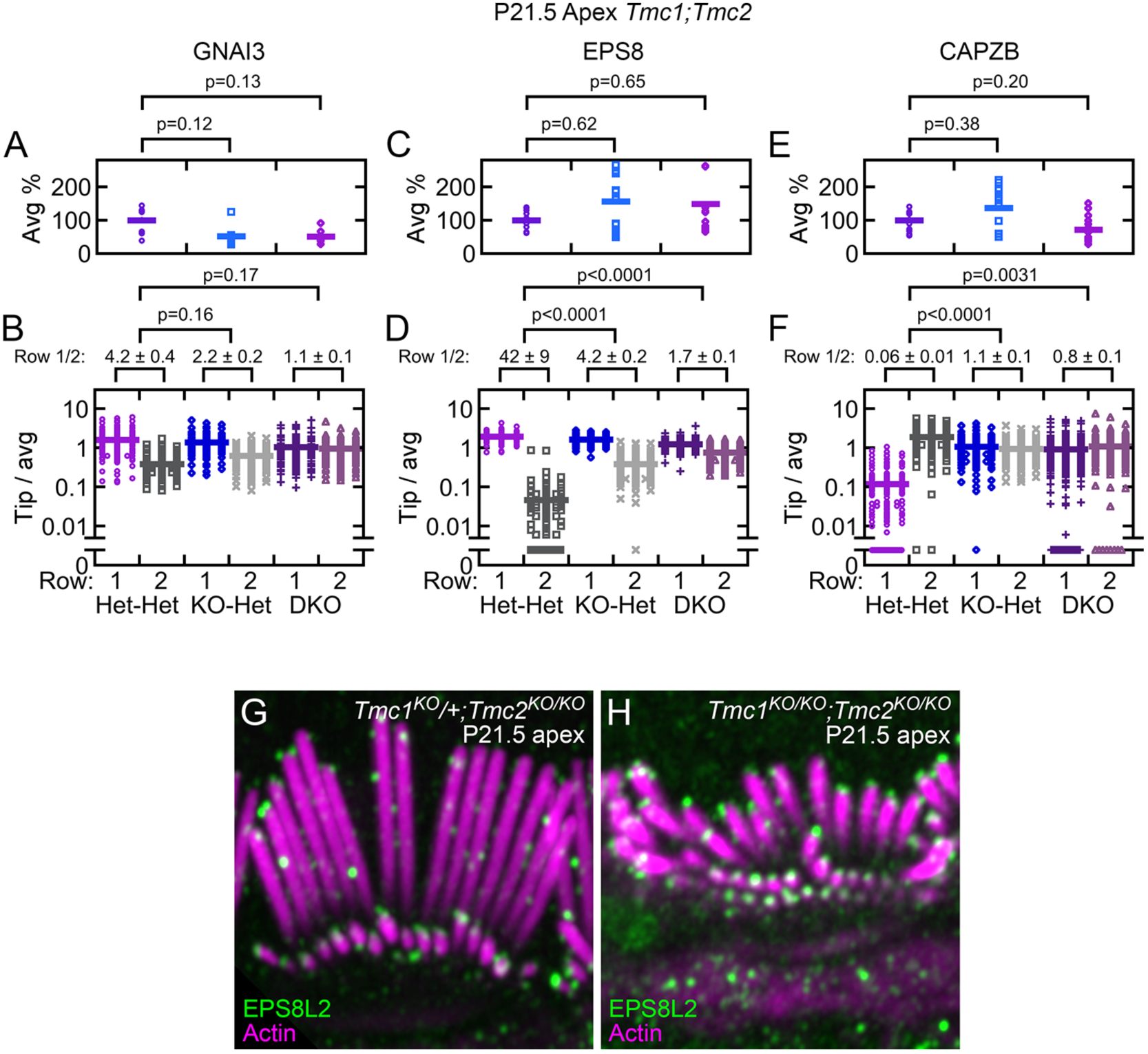
Row 1 and 2 proteins in P21.5 mutant mice. ***A-F***, Measurements of average signal (A, C, E) and row over row average (B, D, F) for GNAI3 (A-B), EPS8 (C-D), and CAPZB (E-F). ***G-H***, EPS8L2 in P21.5 Tmc DKO IHCs. Note labeling at all tips in *Tmc* DKO. Left panel (G), *Tmc1*^*KO/KO*^; *Tmc2*^*KO*^/+ control apical IHCs. Right panel (H), *Tmc1*^*KO/KO*^; *Tmc2*^*KO/KO*^ DKO apical IHCs. Control and KO panels were from littermates, with identical sample preparation and confocal imaging parameters. The gain of the antibody signal was adjusted linearly to reveal the staining pattern in each control sample, and the corresponding KO sample used the same gain settings. Panels are 12 µm wide.

## Supplemental Table

**Table S1.** Antibodies used in this study.

| Antigen | Anti-body name | Antigen | Dilution | Validation | Source | RRID |
| --- | --- | --- | --- | --- | --- | --- |
| GNAI3 | Anti-GNAI3 | C-terminal mouse GNAI3 peptide (KNNLKECGLY) | 1:500 | [3] | Sigma G4040 | RRID: AB_259896 |
| GPSM2 | Anti-GPSM2 | Human GPSM2 amino acids 353-495 | 1:250 | n/a | Sigma HPA007327 | RRID: AB_1849941 |
| EPS8 | Clone 15 | Mouse EPS8 aa 628-821 | 1:250 | [4] | BD Bioscience 610143 | RRID: AB_397544 |
| MYO15A-S | TF1 | Mouse MYO15A-S amino acids 2157–2347 | 1:250 | [5, 6]; this study | Thomas Friedman lab | n/a |
| MYO15A-L | PB888 | Mouse MYO15A-L amino acids 981-998 | 1:250 | [7] | Thomas Friedman lab | n/a |
| Pan-MYO15A | PB48 | Mouse MYO15A-S amino acids 2380–2402 | 1:250 | [5, 6] | Thomas Friedman lab | n/a |
| WHRN | 8127 | C-terminal mouse WHRN peptide with added Cys (KERDYIDFLVTEFN VML-C) | 1:250 | WHRN cDNA expression in cells | Ulrich Müller lab | n/a |
| CAPZB | Anti-CAPZB | C-terminal peptide of human CAPZB2 | 1:250 | [1] | Millipore AB6017 | RRID: AB_10806205 |
| EPS8L2 | Anti-EPS8L2 | Human EPS8L2 amino acids 615-715 | 1:250 | [8] | Abcam ab57571 | RRID: AB_941458 |
| TWF2 | Anti-TWF2 | TWF2>TWF1 | 1:100 | *Twf2*<sup>KO/KO</sup> mice; this study | Stefan Heller lab | n/a |

## References

1. Fettiplace R., and Kim K.X. (2014). The physiology of mechanoelectrical transduction channels in hearing. Physiol Rev. 94, 951–986.

2. Barr-Gillespie P.G. (2015). Assembly of hair bundles, an amazing problem for cell biology. Mol Biol Cell. 26, 2727–2732.

3. Tilney L.G., Tilney M.S., and DeRosier D.J. (1992). Actin filaments, stereocilia, and hair cells: how cells count and measure. Ann. Rev. Cell Biol. 8, 257–274.

4. Kaltenbach J.A., Falzarano P.R., and Simpson T.H. (1994). Postnatal development of the hamster cochlea. II. Growth and differentiation of stereocilia bundles. J Comp Neurol. 350, 187–198.

5. Denman-Johnson K., and Forge A. (1999). Establishment of hair bundle polarity and orientation in the developing vestibular system of the mouse. J Neurocytol. 28, 821–835.

6. Zampini V., Ruttiger L., Johnson S.L., Franz C., Furness D.N., Waldhaus J., Xiong H., Hackney C.M., Holley M.C., Offenhauser N. et al. (2011). Eps8 regulates hair bundle length and functional maturation of mammalian auditory hair cells. PLoS Biol. 9, e1001048.

7. Manor U., Disanza A., Grati M., Andrade L., Lin H., Di Fiore P.P., Scita G., and Kachar B. (2011). Regulation of stereocilia length by myosin XVa and whirlin depends on the actin-regulatory protein Eps8. Curr Biol. 21, 167–172.

8. Fang Q., Indzhykulian A.A., Mustapha M., Riordan G.P., Dolan D.F., Friedman T.B., Belyantseva I.A., Frolenkov G.I., Camper S.A., and Bird J.E. (2015). The 133-kDa N-terminal domain enables myosin 15 to maintain mechanotransducing stereocilia and is essential for hearing. Elife. 4,

9. Belyantseva I.A., Boger E.T., Naz S., Frolenkov G.I., Sellers J.R., Ahmed Z.M., Griffith A.J., and Friedman T.B. (2005). Myosin-XVa is required for tip localization of whirlin and differential elongation of hair-cell stereocilia. Nat Cell Biol. 7, 148–156.

10. Tadenev A.L.D., Akturk A., Devanney N., Mathur P.D., Clark A.M., Yang J., and Tarchini B. (2019). GPSM2-GNAI specifies the tallest stereocilia and defines hair bundle row identity. Curr Biol. 29, 921-934.e4.

11. Mauriac S.A., Hien Y.E., Bird J.E., Carvalho S.D., Peyroutou R., Lee S.C., Moreau M.M., Blanc J.M., Geyser A., Medina C. et al. (2017). Defective Gpsm2/Gαi3 signalling disrupts stereocilia development and growth cone actin dynamics in Chudley-McCullough syndrome. Nat Commun. 8, 14907.

12. Probst F.J., Fridell R.A., Raphael Y., Saunders T.L., Wang A., Liang Y., Morell R.J., Touchman J.W., Lyons R.H., Noben-Trauth K. et al. (1998). Correction of deafness in shaker-2 mice by an unconventional myosin in a BAC transgene. Science. 280, 1444–1447.

13. Mburu P., Mustapha M., Varela A., Weil D., El-Amraoui A., Holme R.H., Rump A., Hardisty R.E., Blanchard S., Coimbra R.S. et al. (2003). Defects in whirlin, a PDZ domain molecule involved in stereocilia elongation, cause deafness in the whirler mouse and families with DFNB31. Nat Genet. 34, 421–428.

14. Tarchini B., Tadenev A.L., Devanney N., and Cayouette M. (2016). A link between planar polarity and staircase-like bundle architecture in hair cells. Development. 143, 3926–3932.

15. Avenarius M.R., Krey J.F., Dumont R.A., Morgan C.P., Benson C.B., Vijayakumar S., Cunningham C.L., Scheffer D.I., Corey D.P., Müller U. et al. (2017). Heterodimeric capping protein is required for stereocilia length and width regulation. J Cell Biol. 216, 3861–3881.

16. Furness D.N., Johnson S.L., Manor U., Ruttiger L., Tocchetti A., Offenhauser N., Olt J., Goodyear R.J., Vijayakumar S., Dai Y. et al. (2013). Progressive hearing loss and gradual deterioration of sensory hair bundles in the ears of mice lacking the actin-binding protein Eps8L2. Proc Natl Acad Sci U S A. 110, 13898–13903.

17. Waguespack J., Salles F.T., Kachar B., and Ricci A.J. (2007). Stepwise morphological and functional maturation of mechanotransduction in rat outer hair cells. J Neurosci. 27, 13890–13902.

18. Lelli A., Asai Y., Forge A., Holt J.R., and Geleoc G.S. (2009). Tonotopic gradient in the developmental acquisition of sensory transduction in outer hair cells of the mouse cochlea. J Neurophysiol. 101, 2961–2973.

19. Beurg M., Cui R., Goldring A.C., Ebrahim S., Fettiplace R., and Kachar B. (2018). Variable number of TMC1-dependent mechanotransducer channels underlie tonotopic conductance gradients in the cochlea. Nat Commun. 9, 2185.

20. Kawashima Y., Geleoc G.S., Kurima K., Labay V., Lelli A., Asai Y., Makishima T., Wu D.K., Della Santina C.C., Holt J.R. et al. (2011). Mechanotransduction in mouse inner ear hair cells requires transmembrane channel-like genes. J Clin Invest. 121, 4796–4809.

21. Pan B., Geleoc G.S., Asai Y., Horwitz G.C., Kurima K., Ishikawa K., Kawashima Y., Griffith A.J., and Holt J.R. (2013). TMC1 and TMC2 are components of the mechanotransduction channel in hair cells of the mammalian inner ear. Neuron. 79, 504–515.

22. Pan B., Akyuz N., Liu X.-P., Asai Y., Nist-Lund C., Kurima K., Derfler B.H., György B., Limapichat W., Walujkar S. et al. (2018). TMC1 forms the pore of mechanosensory transduction channels in vertebrate inner ear hair cells. Neuron. in press.

23. Pacentine I.V., and Nicolson T. (2019). Subunits of the mechano-electrical transduction channel, Tmc1/2b, require Tmie to localize in zebrafish sensory hair cells. PLoS Genet. 15, e1007635.

24. Zhao B., Wu Z., Grillet N., Yan L., Xiong W., Harkins-Perry S., and Muller U. (2014). TMIE is an essential component of the mechanotransduction machinery of cochlear hair cells. Neuron. 84, 954–967.

25. Caberlotto E., Michel V., Foucher I., Bahloul A., Goodyear R.J., Pepermans E., Michalski N., Perfettini I., Alegria-Prevot O., Chardenoux S. et al. (2011). Usher type 1G protein sans is a critical component of the tip-link complex, a structure controlling actin polymerization in stereocilia. Proc Natl Acad Sci U S A. 108, 5825–5830.

26. Vélez-Ortega A.C., Freeman M.J., Indzhykulian A.A., Grossheim J.M., and Frolenkov G.I. (2017). Mechanotransduction current is essential for stability of the transducing stereocilia in mammalian auditory hair cells. Elife. 6,

27. Stepanyan R., and Frolenkov G.I. (2009). Fast adaptation and Ca2+ sensitivity of the mechanotransducer require myosin-XVa in inner but not outer cochlear hair cells. J Neurosci. 29, 4023–4034.

28. Delprat B., Michel V., Goodyear R., Yamasaki Y., Michalski N., El-Amraoui A., Perfettini I., Legrain P., Richardson G., Hardelin J.P. et al. (2005). Myosin XVa and whirlin, two deafness gene products required for hair bundle growth, are located at the stereocilia tips and interact directly. Hum Mol Genet. 14, 401–410.

29. Mathur P.D., Zou J., Zheng T., Almishaal A., Wang Y., Chen Q., Wang L., Vashist D., Brown S., Park A. et al. (2015). Distinct expression and function of whirlin isoforms in the inner ear and retina: an insight into pathogenesis of USH2D and DFNB31. Hum Mol Genet. 24, 6213–6228.

30. Liang Y., Wang A., Belyantseva I.A., Anderson D.W., Probst F.J., Barber T.D., Miller W., Touchman J.W., Jin L., Sullivan S.L. et al. (1999). Characterization of the human and mouse unconventional myosin XV genes responsible for hereditary deafness DFNB3 and shaker 2. Genomics. 61, 243–258.

31. Lelli A., Michel V., Boutet de Monvel J., Cortese M., Bosch-Grau M., Aghaie A., Perfettini I., Dupont T., Avan P., El-Amraoui A. et al. (2016). Class III myosins shape the auditory hair bundles by limiting microvilli and stereocilia growth. J Cell Biol. 212, 231–244.

32. Ebrahim S., Avenarius M.R., Grati M., Krey J.F., Windsor A.M., Sousa A.D., Ballesteros A., Cui R., Millis B.A., Salles F.T. et al. (2016). Stereocilia-staircase spacing is influenced by myosin III motors and their cargos espin-1 and espin-like. Nat Commun. 7, 10833.

33. Glowatzki E., Ruppersberg J.P., Zenner H.P., and Rusch A. (1997). Mechanically and ATP-induced currents of mouse outer hair cells are independent and differentially blocked by d-tubocurarine. Neuropharmacology. 36, 1269–1275.

34. Gale J.E., Marcotti W., Kennedy H.J., Kros C.J., and Richardson G.P. (2001). FM1-43 dye behaves as a permeant blocker of the hair-cell mechanotransducer channel. J Neurosci. 21, 7013–7025.

35. Meyers J.R., MacDonald R.B., Duggan A., Lenzi D., Standaert D.G., Corwin J.T., and Corey D.P. (2003). Lighting up the senses: FM1-43 loading of sensory cells through nonselective ion channels. J Neurosci. 23, 4054–4065.

36. Perrin B.J., Sonnemann K.J., and Ervasti J.M. (2010). β-actin and γ-actin are each dispensable for auditory hair cell development but required for stereocilia maintenance. PLoS Genet. 6, e1001158.

37. Tilney L.G., Tilney M.S., and Cotanche D.A. (1988). Actin filaments, stereocilia, and hair cells of the bird cochlea. V. How the staircase pattern of stereociliary lengths is generated. J. Cell Biol. 106, 355–365.

38. Tompkins N., Spinelli K.J., Choi D., and Barr-Gillespie P.G. (2017). A model for link pruning to establish correctly polarized and oriented tip links in hair bundles. Biophys J. 113, 1868–1881.

39. Offenhauser N., Borgonovo A., Disanza A., Romano P., Ponzanelli I., Iannolo G., Di Fiore P.P., and Scita G. (2004). The eps8 family of proteins links growth factor stimulation to actin reorganization generating functional redundancy in the Ras/Rac pathway. Mol Biol Cell. 15, 91–98.

40. Bird J.E., Takagi Y., Billington N., Strub M.P., Sellers J.R., and Friedman T.B. (2014). Chaperone- enhanced purification of unconventional myosin 15, a molecular motor specialized for stereocilia protein trafficking. Proc Natl Acad Sci U S A. 111, 12390–12395.

41. Giese A.P.J., Tang Y.Q., Sinha G.P., Bowl M.R., Goldring A.C., Parker A., Freeman M.J., Brown S.D.M., Riazuddin S., Fettiplace R. et al. (2017). CIB2 interacts with TMC1 and TMC2 and is essential for mechanotransduction in auditory hair cells. Nat Commun. 8, 43.

42. Goodyear R.J., Marcotti W., Kros C.J., and Richardson G.P. (2005). Development and properties of stereociliary link types in hair cells of the mouse cochlea. J Comp Neurol. 485, 75–85.

43. Rzadzinska A.K., Nevalainen E.M., Prosser H.M., Lappalainen P., and Steel K.P. (2009). MyosinVIIa interacts with Twinfilin-2 at the tips of mechanosensory stereocilia in the inner ear. PLoS One. 4, e7097.

44. Peng A.W., Belyantseva I.A., Hsu P.D., Friedman T.B., and Heller S. (2009). Twinfilin 2 regulates actin filament lengths in cochlear stereocilia. J Neurosci. 29, 15083–15088.

45. Spinelli K.J., and Gillespie P.G. (2009). Bottoms up: transduction channels at tip link bases. Nat Neurosci. 12, 529–530.

46. Assad J.A., and Corey D.P. (1992). An active motor model for adaptation by vertebrate hair cells. J. Neurosci. 12, 3291–3309.

47. Hackney C.M., Mahendrasingam S., Penn A., and Fettiplace R. (2005). The concentrations of calcium buffering proteins in mammalian cochlear hair cells. J Neurosci. 25, 7867–7875.

48. Dumont R.A., Lins U., Filoteo A.G., Penniston J.T., Kachar B., and Gillespie P.G. (2001). Plasma membrane Ca2+-ATPase isoform 2a is the PMCA of hair bundles. J. Neurosci. 21, 5066–5078.

49. Chen Q., Mahendrasingam S., Tickle J.A., Hackney C.M., Furness D.N., and Fettiplace R. (2012). The development, distribution and density of the plasma membrane calcium ATPase 2 calcium pump in rat cochlear hair cells. Eur J Neurosci. 36, 2302–2310.

50. Kuznetsova A., Brockhoff PB, and Rhb C. (2017). lmerTest Package: Tests in Linear Mixed Effects Models. Journal of Statistical Software. 82, 1–26.

51. Narayan K., and Subramaniam S. (2015). Focused ion beams in biology. Nat Methods. 12, 1021–1031.

## Supplemental References

1. Avenarius M.R., Krey J.F., Dumont R.A., Morgan C.P., Benson C.B., Vijayakumar S., Cunningham C.L., Scheffer D.I., Corey D.P., Müller U. et al. (2017). Heterodimeric capping protein is required for stereocilia length and width regulation. J Cell Biol. 216, 3861–3881.

2. Krey J.F., Sherman N.E., Jeffery E.D., Choi D., and Barr-Gillespie P.G. (2015). The proteome of mouse vestibular hair bundles over development. Sci Data. 2, 150047.

3. Leopoldt D., Harteneck C., and Nürnberg B. (1997). G proteins endogenously expressed in Sf 9 cells: interactions with mammalian histamine receptors. Naunyn Schmiedebergs Arch Pharmacol. 356, 216–224.

4. Zampini V., Ruttiger L., Johnson S.L., Franz C., Furness D.N., Waldhaus J., Xiong H., Hackney C.M., Holley M.C., Offenhauser N. et al. (2011). Eps8 regulates hair bundle length and functional maturation of mammalian auditory hair cells. PLoS Biol. 9, e1001048.

5. Liang Y., Wang A., Belyantseva I.A., Anderson D.W., Probst F.J., Barber T.D., Miller W., Touchman J.W., Jin L., Sullivan S.L. et al. (1999). Characterization of the human and mouse unconventional myosin XV genes responsible for hereditary deafness DFNB3 and shaker 2. Genomics. 61, 243–258.

6. Belyantseva I.A., Boger E.T., and Friedman T.B. (2003). Myosin XVa localizes to the tips of inner ear sensory cell stereocilia and is essential for staircase formation of the hair bundle. Proc. Natl. Acad. Sci. USA. 100, 13958–13963.

7. Fang Q., Indzhykulian A.A., Mustapha M., Riordan G.P., Dolan D.F., Friedman T.B., Belyantseva I.A., Frolenkov G.I., Camper S.A., and Bird J.E. (2015). The 133-kDa N-terminal domain enables myosin 15 to maintain mechanotransducing stereocilia and is essential for hearing. Elife. 4,

8. Furness D.N., Johnson S.L., Manor U., Ruttiger L., Tocchetti A., Offenhauser N., Olt J., Goodyear R.J., Vijayakumar S., Dai Y. et al. (2013). Progressive hearing loss and gradual deterioration of sensory hair bundles in the ears of mice lacking the actin-binding protein Eps8L2. Proc Natl Acad Sci U S A. 110, 13898–13903.

